# Characterizing the genetic basis of copper toxicity in *Drosophila* reveals a complex pattern of allelic, regulatory, and behavioral variation

**DOI:** 10.1101/2020.05.13.094524

**Authors:** Elizabeth R. Everman, Kristen M. Cloud-Richardson, Stuart J. Macdonald

**Affiliations:** Department of Molecular Biosciences, University of Kansas, 1200 Sunnyside Avenue, Lawrence, KS 66045, USA; Center for Computational Biology, University of Kansas, 2030 Becker Drive, Lawrence, KS 66047

**Keywords:** Heavy metal resistance, copper, life stage-specific, DSPR, QTL, MMP

## Abstract

A range of heavy metals are required for normal cell function and homeostasis. Equally, the anthropogenic release of heavy metals into soil and water sources presents a pervasive health threat. Copper is one such metal; it functions as a critical enzymatic cofactor, but at high concentrations is toxic, and can lead to the production of reactive oxygen species. Using a combination of quantitative trait locus (QTL) mapping and RNA sequencing in the *Drosophila* Synthetic Population Resource (DSPR), we demonstrate that resistance to the toxic effects of ingested copper in *D. melanogaster* is genetically complex, and influenced by allelic and expression variation at multiple loci. Additionally, we find that copper resistance is impacted by variation in behavioral avoidance of copper and may be subject to life-stage specific regulation. Multiple genes with known copper-specific functions, as well as genes that are involved in the regulation of other heavy metals were identified as potential candidates to contribute to variation in adult copper resistance. We demonstrate that nine of 16 candidates tested by RNAi knockdown influence adult copper resistance, a number of which may have pleiotropic effects since they have previously been shown to impact the response to other metals. Our work provides new understanding of the genetic complexity of copper resistance, highlighting the diverse mechanisms through which copper pollution can negatively impact organisms. Additionally, we further support the similarities between copper metabolism and that of other essential and nonessential heavy metals.

## Introduction

Anthropogenic release of heavy metals into soil and water sources presents a pervasive threat with long-lasting ecological, health, and economic impacts (Wu et al. 1975; Wuana and Okieimen 2011; Babin-Fenske and Anand 2011; Gall et al. 2015). Elevated heavy metals have been reported in dozens of organisms at all levels of the ecosystem (Neuberger et al. 1990; Georgieva et al. 2002; Sánchez-Chardi and Nadal 2007; Usmani 2011; Gall et al. 2015; Wright et al. 2015; Plessl et al. 2017; Ecke et al. 2017; Ilunga Kabeya et al. 2018), demonstrating that the toxic effects of heavy metal pollution are wide reaching, and can spread through food webs (Gall et al. 2015; Ilunga Kabeya et al. 2018). Although required for normal physiological function at low concentrations, copper is one of many common environmental heavy metal pollutants linked to mining (Ramirez et al. 2005; Wuana and Okieimen 2011; Wright et al. 2015), pipes used to provide drinking water (Harvey et al. 2016), and pesticide use (de Oliveira-Filho et al. 2004). In its essential role, copper helps bind oxygen, catalyzes enzymatic reactions, and promotes normal neurological development (Hart et al. 1928; Danks 1988; World Health Organization et al. 1996; Uriu-Adams and Keen 2005; Norgate et al. 2006; Navarro and Schneuwly 2017). However, excessive copper exposure ultimately leads to the accumulation of reactive oxygen species, which can cause cellular damage through oxidative stress (Uriu-Adams and Keen 2005; Tchounwou et al. 2008).

Evolutionarily conserved metal-responsive transcription factor 1 (MTF-1) and metallothionein (MT) proteins function as a first line of defense against toxic effects of excessive copper exposure in diverse organisms including humans, flies, fungi, and plants (Macnair 1993; Goldsbrough 2000; Bellion et al. 2007; Calap-Quintana et al. 2017). MTF-1 binds to metal responsive elements of MT genes, increasing MT abundance in copper accumulating cells and allowing excess heavy metal ions to be sequestered until they are removed from the system (Filshie et al. 1971; Stuart et al. 1985; Mokdad et al. 1987; Egli et al. 2003; Balamurugan et al. 2004; Southon et al. 2004). Metal chaperone and transporter proteins such as *Ccs* (Culotta et al. 1997), *Atox1* (Southon et al. 2004; Hatori and Lutsenko 2013), *ATP7* (Norgate et al. 2006), and *CTR1* family transporters (Petris et al. 2003; Guo et al. 2004; Balamurugan et al. 2007; Turski and Thiele 2007; Calap-Quintana et al. 2017; Navarro and Schneuwly 2017) play a crucial role in the response to heavy metal toxicity as well (Egli et al. 2003; Petris et al. 2003; Yepiskoposyan et al. 2006; Janssens et al. 2009). For example, in *Drosophila melanogaster*, high copper exposure decreases translation of *Ctr1A* and *Ctr1B* via *MTF*-*1* to reduce influx of copper ions (Balamurugan et al. 2007; Turski and Thiele 2007; Calap-Quintana et al. 2017; Navarro and Schneuwly 2017), whereas high copper exposure in humans leads to degradation of the *hCTR1* protein (the human ortholog of *Ctr1A/B*) and a reduction in the intracellular concentration of copper (Petris et al. 2003; Guo et al. 2004).

Much of our understanding of the response to copper stress has come from studies that use genetic manipulation to define the roles of metal responsive genes (e.g. Egli et al. 2003, 2006a; Bellion et al. 2007; Kirby et al. 2008; Bahadorani et al. 2010). However, QTL (quantitative trait locus) and GWA (genome-wide association) studies have demonstrated the genetic complexity of the response to heavy metal stress (Willems et al. 2007; Courbot et al. 2007; Zhou et al. 2017). For example, QTL mapping using *Caenorhabditis elegans* recombinant inbred advanced intercross lines (RIAILs) showed that several regions of the genome are involved in the response to cadmium, copper, and silver exposure (Evans et al. 2018). GWA with the *Drosophila* Genetic Reference Panel (DGRP) revealed multiple candidate loci associated with the response to cadmium and lead stress (Zhou et al. 2017). QTL mapping of metal resistance in plants has further demonstrated the role that allelic variation plays in the response to heavy metal stress despite strong selection pressure against metal-sensitive alleles in natural populations (Willems et al. 2007; Courbot et al. 2007; Turner et al. 2010; Arnold et al. 2016). For instance, an interspecific QTL study of two closely related species of *Arabidopsis* (metal-tolerant *A. halleri* x metal-sensitive *A. lyrata petraea*) identified multiple regions of the genome that contributed to zinc and cadmium resistance, and demonstrated that metal resistant alleles had become fixed in the metal tolerant species in the populations sampled (Willems et al. 2007; Courbot et al. 2007). Similarly, sequencing of *A. arenosa* populations locally adapted to serpentine soils revealed strong selection for introgressed alleles from the more tolerant *A. lyrata* (Arnold et al. 2016). These and other examples from *Mimulus gutattus* growing in old copper mine tailings (Allen 1971; Macnair 1983; MacNair et al. 1993; Wright et al. 2015; Selby and Willis 2018) highlight the utility of using a quantitative genomics approach with powerful mapping panels to examine the influence of allelic variation on metal tolerance.

*D. melanogaster* is an important model for understanding the mechanisms involved in the response to toxic heavy metal exposure due to the ease with which it can be genetically manipulated (e.g. Egli et al. 2006a; reviewed in Navarro and Schneuwly 2017) and to the extensive conservation of genes involved in the response to metal ions between flies and humans (Zhou et al. 2017). In addition, the existence of large *Drosophila* mapping panels makes this model system especially well-suited for examining the effect of naturally-occurring alleles on the response to heavy metal stress. In this study, we used the *Drosophila* Synthetic Population Resource (DSPR) (King et al. 2012a; b) to investigate the influence of allelic and expression variation on the response to toxic copper exposure through QTL mapping with more than 1500 strains, and RNA sequencing of copper-resistant and copper-sensitive strains. Because several genes, including *MTF-1* and MTs, respond to multiple heavy metals (e.g. copper, zinc, lead, and cadmium (Calap-Quintana et al. 2017)), and because other’s work has suggested that pleiotropic QTL can underlie genetic variation for multiple metals (Evans et al. 2018), our focus on the genetic architecture of copper resistance has the potential to provide insight into a broader set of genes with allelic and expression variation that influence the response to heavy metal stress.

In this study, we mapped 12 QTL associated with variation in adult female copper resistance, and the implicated genomic regions include genes with known copper-specific functions. Of particular note are *Ccs*, which shuttles copper ions to superoxide dismutase 1 (SOD1) (Schmidt et al. 2000), *Sod3*, which binds extracellular copper ions and is involved in the response to oxidative stress (Blackney et al. 2014), and *CG11825*, a gene that has been previously linked to copper exposure in other studies (Norgate et al. 2007). Copper resistance QTLs include genes with functions linked to other metals, including *Catsup* (zinc, (Navarro and Schneuwly 2017)) and *whd* (lead and cadmium (Strub et al. 2008)), providing new evidence that these genes may also be involved in the response to copper stress. Gene expression differences between a subset of resistant and sensitive DSPR strains additionally showed that many other genes are involved in the response to copper stress. We showed that copper sensitive strains were characterized by increased expression of genes related to mitochondrial function and energy synthesis relative to resistant strains. We tested 16 candidate genes that were implicated by QTL or differential expression analysis using RNAi knockdown in the whole animal and in the anterior region of the midgut. Knockdown of 9 of these genes, including both QTL and RNAseq candidates, influenced copper resistance. Overall, because several of our candidates have not been previously linked to copper stress, we were able to demonstrate that both genes with known copper specific function (such as *Ccs*) and genes that may influence regulation of other heavy metals (such as *trpl*) influence resistance to copper stress. Together, our study provides new insight into the diverse set of genes that impact copper resistance through allelic and expression variation.

## Materials and Methods

### Mapping panel

We reared and phenotyped the >1500 recombinant inbred lines (RILs) that comprise the DSPR (*Drosophila* Synthetic Population Resource) to measure variation in susceptibility to copper stress. The DSPR is a multiparental, advanced generation intercross mapping panel derived from 15 unique and fully-sequenced founder strains, which represent a global sampling of genetic diversity in D. melanogaster. The DSPR consists of two mapping panels (A and B), which are composed of two subpanels (A1 and A2, and B1 and B2). The subpanels were started from the same set of founders, but were maintained independently (see King et al. 2012b for additional details on the mapping panel).

### Rearing and assay conditions

Strains from the DSPR were maintained, reared, and tested in the same incubator under a 12:12hr light:dark photoperiod at 25°C and 50% humidity. To obtain female flies for the adult copper response assay, RNA sequencing, and RNAi validation, adults were transferred to cornmeal-molasses-yeast food, allowed to oviposit for two days, then discarded. Experimental female, presumably mated, flies from the following generation were sorted over CO_2_ and were placed into vials with new cornmeal-molasses-yeast media for 24 hours prior to the start of each assay before they were transferred to copper-supplemented food. All adult assays were performed on 3-5 day old individuals.

### Adult female copper resistance

The adult female response to copper stress was measured as percent survival on media containing 50mM CuSO_4_ following a 48-hour exposure period. As essentially no flies die on control food throughout the span of our assay (Highfill et al. 2016), we did not assess adult female survival on control food in this study. Experimental females were transferred without CO_2_ anesthesia to vials containing 1.8g potato-based Instant *Drosophila* Medium (Carolina Biological Supply Company 173200) hydrated with 8mL 50mM CuSO_4_ (Copper(II) sulfate, Sigma-Aldrich C1297). Instant *Drosophila* Medium is estimated to contain approximately 0.02mM Cu prior to hydration (Maroni and Watson 1985). Copper resistance was measured in a total of 11 batches across the A (N strains = 767) and B (N strains = 789) mapping panels of the DSPR. Each strain was measured in a single batch with 3 vial replicates each containing between 7 and 20 individuals (average number of flies per vial replicate = 19.4). The effect of copper on survival was reported as mean percent survival per strain across the three replicate vials. Retaining vials with fewer than 15 flies did not impact our QTL mapping results in a meaningful way (see below). Hereafter, the adult survival response to 48hr 50mM CuSO_4_ is referred to as adult copper resistance.

### Adult female feeding response to copper-supplemented media

A subset of strains evenly sampled throughout the distribution of B2 subpanel adult copper resistance (0% ± 0 S.E. – 98.4% ± 1.59 S.E.) were used to measure the effect of copper exposure on food intake. We measured food intake in 3 blocks with at least two vial replicates of 20 females per strain (N = 95) per treatment (control vs. copper), following a protocol modified from (Shell et al. 2018).

Briefly, we added 1% Erioglaucine Disodium Salt (Sigma-Aldrich 861146), a blue dye, to water and to 50mM CuSO_4_ no more than 24 hours prior to the assay to avoid dye decomposition. We hydrated 0.9g Instant *Drosophila* Medium with 4mL liquid, and flies were allowed to consume dyed food for 24 hours before they were frozen for up to 5 hours. No flies died during the 24-hour period. Subsequently, flies were homogenized with 3-4 glass beads in 600μL distilled water for 45 seconds using a Mini-Beadbeater-96 (BioSpec Products). Homogenate was centrifuged for 5 minutes at 14,000rpm, and 200μL supernatant was transferred to a 96-well plate. Fly homogenate was frozen for up to 48 hours before absorbance at 630nm was measured with a BioTek Multimode Microplate reader (Synergy 2 v.1.03). Two water blanks and 14 standards ranging from 6.25 x 10^-5^% to 0.006% dye in water were prepared fresh for each block, and were included in each plate to determine the dye concentration of fly homogenate, and to assess consistency among blocks. Absorbance readings for standards were highly correlated across plates and blocks (Table S1). To calculate the estimated amount of dye consumed, we used a linear model to find the slope and intercept of the standard curve (Concentration of Standard ∼ Absorbance * Block). Estimated percent dye in each fly homogenate sample was determined with the equation

**Table 1.** Summary of QTL identified for response to copper stress by panel and life stage.

| **Adult Copper Resistance QTL: A panel** |  |  |  |  |  |  |
| --- | --- | --- | --- | --- | --- | --- |
| **QTL** | **Peak LOD** | **Chr** | **Physical Interval (Mb)<sup>a</sup>** | **Genetic Interval (cM)** | **Variance Explained** | **No. Genes<sup>b</sup>** |
| Q1 | 10.13 | 2L | 14.22-15.07 | 49.52-60.62 | 5.91 | 73 |
| Q2 | 9.76 | 2L | 15.77-16.38 | 51.35-51.88 | 5.70 | 68 |
| Q3A | 11.3 | 2L | 18.87-20.09 | 53.39-53.85 | 6.56 | 154 |
| Q6 | 8.01 | 2R | 7.03-7.72 | 63.56-64.89 | 4.70 | 108 |
| Q7 | 8.54 | 2R | 18.54-19.03 | 100.24-102.17 | 5.01 | 92 |
| Q10 | 6.88 | 3R | 2.48-2.95 | 47.56-47.74 | 4.05 | 40 |
| Q12 | 7.24 | 3R | 21.01-21.46 | 86.55-88.14 | 4.26 | 78 |

| **Adult Copper Resistance QTL: B panel** |  |  |  |  |  |  |
| --- | --- | --- | --- | --- | --- | --- |
| **QTL** | **Peak LOD** | **Chr** | **Physical Interval (Mb)<sup>a</sup>** | **Genetic Interval (cM)** | **Variance Explained** | **No. Genes<sup>b</sup>** |
| Q3B | 9.21 | 2L | 17.95-20.31 | 52.92-53.90 | 5.24 | 259 |
| Q4 | 9.44 | 2R | 1.41-3.69 | 54.99-57.07 | 5.36 | 247 |
| Q5 | 8.46 | 2R | 4.08-6.58 | 57.77-62.69 | 4.82 | 353 |
| Q8 | 7.88 | 3L | 15.50-16.82 | 42.90-44.06 | 4.49 | 240 |
| Q9 | 6.97 | 3L | 17.35-19.47 | 44.48-45.78 | 3.99 | 224 |
| Q11 | 7.05 | 3R | 14.57-14.98 | 63.92-65.00 | 4.03 | 41 |

| **Copper-specific Developmental Viability QTL: B panel** |  |  |  |  |  |  |
| --- | --- | --- | --- | --- | --- | --- |
| **QTL** | **Peak LOD** | **Chr** | **Physical Interval (Mb)<sup>a</sup>** | **Genetic Interval (cM)** | **Variance Explained** | **No. Genes<sup>b</sup>** |
| Q13 | 7.11 | 2R | 13.30-13.95 | 82.37-84.50 | 27.9 | 125 |
| Q14 | 7.74 | 2R | 16.19-16.66 | 90.86-92.45 | 30.0 | 60 |
| Q15 | 8.65 | 3R | 14.52-15.06 | 63.79-65.21 | 32.8 | 62 |

<sup>a</sup>Physical intervals are based on FlyBase release 5 of the *D. melanogaster* reference genome.
<sup>b</sup>Protein coding genes only. All genes including ncRNA and pseudogenes are included in Table S4.

*% Dye in Sample = (0.002443 x absorbance) − 0.0001465*

Variation in feeding behavior among DSPR strains on copper and control food was assessed with a two-way ANOVA with an interaction (% Dye Consumed ∼ DSPR Strain * Treatment), and effect size was calculated using Cohen’s F (R package sjstats) (Cohen 1988; Lüdecke 2018). The correlation between average percent dye consumed and average adult copper resistance was assessed with a linear model that included an interaction between percent dye consumed and treatment in addition to the additive effects of these factors.

### Copper-specific developmental response

Developmental viability was estimated in the B panel from 100 strains that were evenly sampled from throughout the distribution of B1 and B2 subpanel adult copper resistance (0% ± 0 S.E. – 98.4% ± 1.59 S.E.). Approximately 100 females per strain were allowed to oviposit on cornmeal-molasses-yeast media for 17-20 hours in 6oz polypropylene *Drosophila* bottles (Genesee Scientific: 32-130) with yeast paste to encourage egg laying. Following oviposition, remaining yeast paste was removed and embryos were gently dislodged from the media surface by rinsing with 1X PBS and swirling with a small, bristled paintbrush. Subsequently, for each strain, we arrayed multiple aliquots of 10μL of embryos suspended in 1X PBS onto a petri dish containing 2% agar dyed blue with Erioglaucine Disodium Salt (Sigma-Aldrich 861146; 8mg/mL).

For each dish we aliquoted eggs into 14 cells (Figure S1), photographed the dish (Nikon D3200, 105mm 1:2.8 DG Sigma Macro lens), and the number of embryos within each cell was recorded with ImageJ (v. 1.51s). Embryos from each cell (30 – 306 embryos, average = 125 embryos) were then transferred with a rubber, bristleless paintbrush to vials containing control or 2mm CuSO_4_ hydrated Instant *Drosophila* Medium (1.8g media plus 8ml liquid). The rubber paintbrush was examined after each egg transfer to ensure all eggs had been transferred to the vial. The developmental response to copper was assessed with 4-7 replicates per treatment for each strain (mean replicates per strain = 6.8). We used a lower copper concentration in this assay because previous reports have shown that the larval life stage is much more susceptible to copper toxicity compared to adults (Bahadorani and Hilliker 2009).

Copper stress has the potential to reduce the number of individuals that complete development from egg to adult as well as the time needed for individuals to complete development. To estimate the effect of copper exposure on development time, for each experimental vial we recorded the number of days between set up and the first emergence of adults. To assess the effect of copper on developmental viability, we calculated the proportion of embryos in each vial that eclosed as adults in the seven days following the day of first emergence. Developmental viability was square root transformed to improve deviation from normality within treatment (Shapiro Wilks Test on transformed data; W = 0.99, P = 0.04). From here forward, square root transformed developmental viability is simply referred to as developmental viability and all subsequent analyses were performed on square root transformed data. Vials were monitored daily for 30 days after set up. Of the 1,356 vials included in this assay, 100 copper treatment vials yielded zero flies. These vials were given a development time of 30 days.

We used a 2-way ANOVA with an interaction to measure the effect of strain and treatment on each developmental response. The DSPR strains we used in this study varied in development time (F_99,579_ = 11.8, P < 0.00001; Table S2A) and developmental viability on control food (F_(99,579)_ = 31.6, P < 0.00001; Table S2B). Furthermore, regression analysis demonstrated that development time and developmental viability in control and copper conditions were correlated (development time: F_(1,98)_ = 61.0, P < 0.0001, R^2^ = 38%; Table S2C, Figure S2A; developmental Viability: F_(1,98)_ = 54.1, P < 0.0001, R^2^ = 36%; Table S2D, Figure S2B). Therefore, we regressed out variation in control development time and control developmental viability with linear models to more directly assess the effect of copper stress on these metrics of development. Residual development time and residual developmental viability are referred to from hereafter as copper-specific development time and copper-specific developmental viability, respectively.

### Heritability

We estimated the genetic and phenotypic variances of adult copper resistance, control and copper feeding responses, copper-specific development time, and copper-specific developmental viability using linear mixed models (lme and varcomp functions in R; (R package: APE Paradis et al. 2004; R package: nlme Pinheiro et al. 2017)). For all responses, panel-specific broad-sense heritabilities were calculated as the proportion of the total strain-specific variation in the response to copper explained by the estimated genetic variance component (Lynch and Walsh 1998).

### QTL mapping of life stage-specific response to copper stress

We used the DSPRqtl package in R (github.com/egking/DSPRqtl; FlyRILs.org), to identify QTL associated with variation in adult copper resistance, adult feeding response on control and on copper-supplemented food, copper-specific development time, and copper-specific developmental viability. QTL mapping was performed for each mapping panel (A and B) and phenotype separately. At each position in the genome, for each strain, we can estimate the additive probability that the segment of the genome was inherited from each of the 8 DSPR founders. QTL were identified by regressing the strain mean phenotype on these probabilities, and significance thresholds were assigned following 1000 permutations of the data (King et al. 2012a; b). For adult copper resistance, peak positions for each QTL were estimated with 2-LOD support intervals (King et al. 2012a). Because fewer strains were used to measure the feeding and copper-specific development traits, we used a 3-LOD drop to determine peak support intervals for these traits (King et al. 2012a). Gene ontology (GO) analysis was performed without normalizing for gene length for genes included in peak intervals for each trait and mapping panel separately (FlyMine.org (Lyne et al. 2007)).

Adult copper resistance varied between the A1 and A2 subpanels but did not vary between the B subpanels (A panel: F_1,2289_ = 12.64; p < 0.001; B panel: F_1,2495_ = 0.03; p = 0.86; Figure S3; Table S2E,F). Therefore, subpanel was included as a model covariate only in the QTL analysis of panel A. Phenotyping batch also significantly influenced variation in adult copper resistance in both the A and B panels (Table S2E,F). However, including batch as a covariate in the QTL mapping model did not drastically alter the estimation of LOD scores for either panel (A panel correlation between LOD scores = 99%; B panel correlation = 98%; Figure S4A), so we only present data from the models lacking batch as a covariate. Because the development assay was conducted on 100 strains across 15 batches, DSPR strain was highly confounded with batch. Therefore, we did not include batch or subpopulation as a covariate in the QTL mapping models for copper-specific development time or copper-specific developmental viability. As each strain assessed in the feeding response assay was measured in each of three blocks, block was not included in the model for either average feeding response on control or copper food.

To determine whether including vials containing relatively few flies influenced QTL mapping results due to mis-estimated phenotype means, we additionally mapped variation in adult copper resistance using only data from vials containing at least 15 flies (removing 316 - 7% - of the vials). LOD scores for the full data set were highly correlated with those for the reduced dataset for each panel (A panel correlation = 99%; B panel correlation = 99%; Figure S4B), so we only present data from analyses using all vials.

### Differential gene expression in high and low adult copper resistance strains

We examined gene expression variation in a subset of 10 strains (6 with high adult resistance: 76 – 98% survival, and 4 with low adult resistance: 0.0 – 18% survival) from the B panel to explore how adult copper resistance class and gene expression interact when individuals are exposed to 50mM CuSO_4_. Twenty experimental females from each DSPR strain were transferred to Instant *Drosophila* Medium hydrated with either water as a control or 50mM CuSO_4_ for 9 hours. No individuals died during the 9-hour exposure period. Following exposure, 10 females from each strain and treatment were flash frozen in liquid nitrogen, placed in TRIzol Reagent (Invitrogen, 15596018), and immediately stored at −80°C. RNA was extracted from each of the 20 samples with the Direct-zol RNA Miniprep kit (Zymo Research, R2050), eluted in 100μL water and stored at −80°C. We prepared libraries with the TruSeq Stranded mRNA kit (Illumina, 20020595), and paired-end 37-bp mRNA libraries were each sequenced to ∼20 million reads on an Illumina NextSeq 550 at the University of Kansas Genome Sequencing Core.

Sequence quality assessment and trimming were performed using fastp (Chen et al. 2018). We used kallisto to perform pseudoalignment-based mapping of reads (Ensembl transcriptome release 90) (Bray et al. 2016), and performed differential expression analysis with sleuth (v.0.30.0) using likelihood ratio tests (Pimentel et al. 2017). Gene expression is likely to vary between the different DSPR strains; however, we were primarily interested in understanding whether there are consistent differences in gene expression between high and low resistance classes of strains. Given this interest, we treated each strain as a biological replicate of the high and low resistance classes and did not include DSPR strain in differential expression models. After determining that the interaction between resistance class and treatment did not influence expression of any gene at a 5% FDR (False Discovery Rate), we tested the additive effects of resistance class and treatment on gene expression, referred to from here as the full model (full model: ∼ TRT + RES vs reduced model: ∼ 1). We also examined the influence of each term independently in two additional models. The effect of treatment alone was assessed by accounting for resistance class (treatment model: ∼ TRT + RES vs. reduced model: ∼ RES), and the effect of resistance class alone was assessed by accounting for treatment (resistance model: ∼ TRT + RES vs. reduced model: ∼ TRT). From here on, these term-specific models are referred to as the treatment model and the resistance model, respectively. Significantly differentially expressed (DE) genes for each model were identified with a 5% FDR threshold.

We generated three lists of significantly differentially expressed (DE) genes: full model DE genes, treatment model DE genes, and resistance model DE genes. Sleuth applies a filter against genes with low expression (Pimentel et al. 2017). We applied an additional filter following sleuth analysis to remove genes from DE gene lists with average expression of less than 1 TPM (Transcripts Per Million). Additionally, we eliminated genes with expression variance greater than or equal to 1 TPM in any of the following four categories: sensitive strains, control treatment; sensitive strains, copper treatment; resistant strains, control treatment; resistant strains, copper treatment (Figure S5). We used principal components analysis (PCA) to examine the effect of treatment and resistance class using quantile normalized TPM data for DE genes. DE gene lists were examined for co-regulated clusters of genes using Clust (Abu-Jamous and Kelly 2018). Gene ontology (GO) analysis was performed for each cluster and for each of the DE gene lists in their entirety (FlyMine.org (Lyne et al. 2007)).

### RNAi knockdown of candidate genes associated with adult copper resistance

QTL mapping and RNAseq of adult females provided several candidate genes that were implicated by one or both of these experiments. TRiP UAS-RNAi strains (Perkins et al. 2015) for candidate genes, as well as a control UAS-LUC.VALIUM10 strain, were obtained from the Bloomington Drosophila Stock Center (BDSC) (Table S3). Crosses involved 10 TRiP males and 10 virgin females from Gal4-expressing driver strains. Each TRiP strain was crossed to both a ubiquitous Gal4-expressing driver strain (BDSC 25374) and an anterior midgut specific Gal4-expressing RNAi driver strain (1099 from Nicholas Buchon, flygut.epfl.ch). Three candidate gene TRiP strains (*swm*, *Catsup*, and *CG11825*) produced too few flies to test when crossed to the ubiquitous Gal4-expressing driver, and were thus excluded from our analysis. We tested two TRiP UAS RNAi strains for the genes *CG5235*, *MtnC*, and *ZnT41F* to assess the consistency in the effect of gene knockdown on copper survival (Table S3). An average of 19.3 Gal4-UAS RNAi females (min 7) were transferred to Instant Drosophila Medium hydrated with 50mM CuSO_4_ using an average of 16.8 (min 10) replicate vials per genotype (a total of 203-365 individuals per genotype) across four batches. We counted flies daily until all were dead, and the response to copper stress in these RNAi knockdown genotypes was quantified as average lifespan. We chose to measure lifespan on copper instead of percent survival at 48 hours (as in our DSPR screen) because we had no meaningful a priori expectation that survival would be variable at 48 hours among the RNAi genotypes, and knockdown in genes hypothesized to influence the response to copper toxicity could drastically reduce or extend survival. To establish that GAL4-UAS-RNAi genotypes were not inherently unhealthy, we additionally placed 20 such females from each cross on Instant *Drosophila* Medium hydrated with water to assess overall viability. No individuals died on copper-free media through the duration of the RNAi assay. We compared copper resistance for each RNAi knockdown to its respective control using per vial average lifespan controlling for batch with a two-way ANOVA (Average Lifespan ∼ Strain * Batch) with planned comparisons. These analyses were performed separately for each GAL4 driver.

### Data availability

All raw data and images generated from this study including adult and developmental copper resistance traits, feeding data, raw QTL mapping data, normalized TPM expression data, and RNAi data are available at FigShare. File S1 contains descriptions for all accompanying data files. DSPR genotype information is publicly available at http://wfitch.bio.uci.edu/~dspr/. RNAseq reads will be submitted to the NCBI SRA prior to acceptance. Unless otherwise stated, all analyses were performed in R (v. 3.6.2) (R Core Team 2017).

## Results

### Abundant variation in adult female copper resistance

We measured adult copper resistance in females from over 1,500 DSPR RILs by exposing close to 60 flies (3 vials of 20 flies) from each strain to 50mM CuSO_4_ for 48 hours. Phenotypic variation and heritability were high for female copper resistance in both the A and B panels of the DSPR (A panel: H^2^ = 83.0%; B panel: H^2^ = 78.8%; Figure 1).

**Figure 1.**
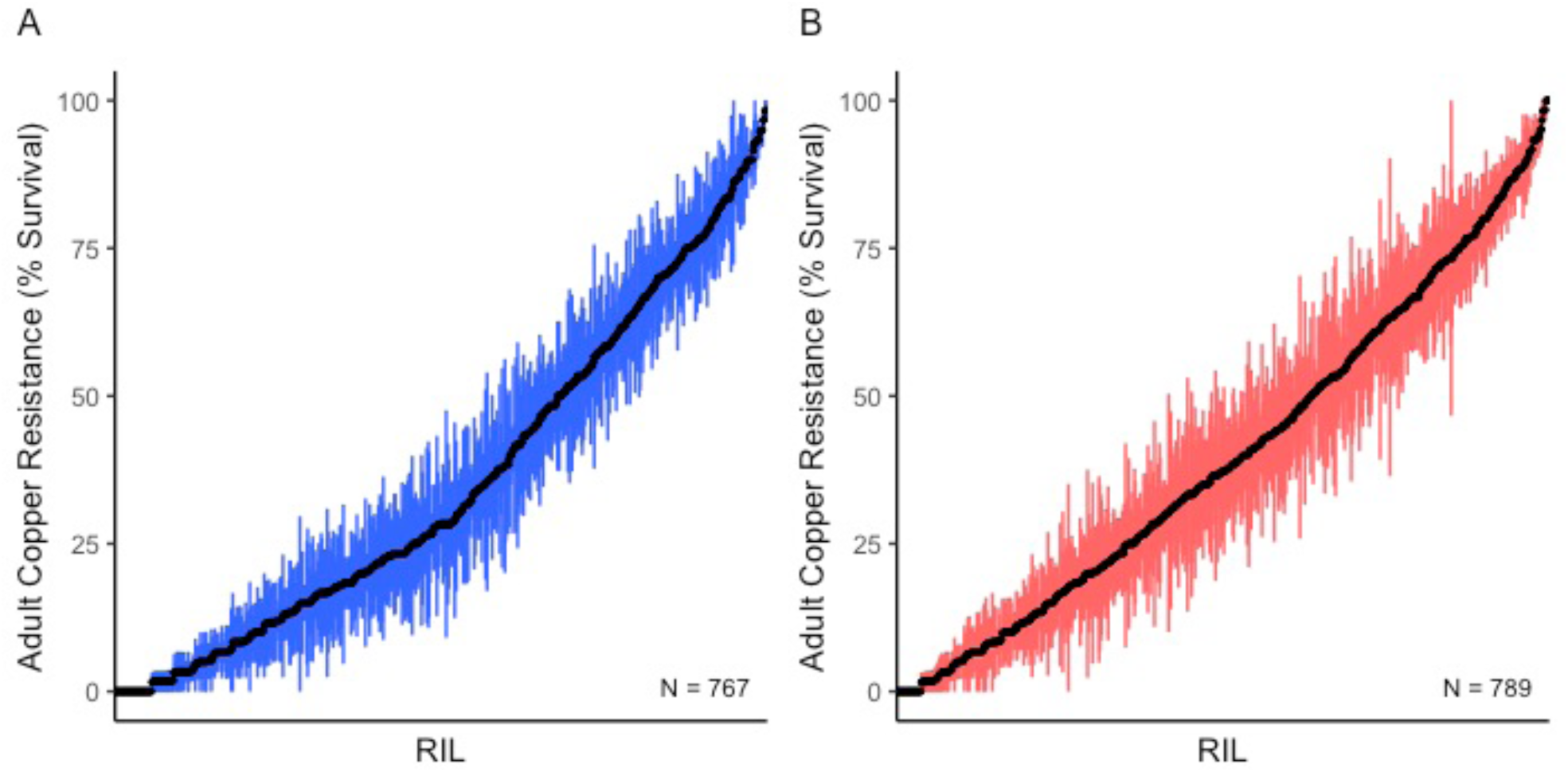
Variation in mean female adult copper resistance (± SE) per DSPR strain in the A (A) and B (B) panels following 48-hour exposure to 50mM CuSO4. Recombinant inbred line (RIL) is ordered by magnitude of response along the x axis.

### Adult female feeding response to copper-supplemented media

Since the toxicity of copper in our assay likely stems from ingestion, we were interested in flies’ feeding response to copper-supplemented media. Using a sample of 95 strains from the B2 subpanel that spanned the distribution of adult copper resistance (from 0% −98.4% survival), we tested the effect of 50mM CuSO_4_ on feeding behavior. We estimated feeding by measuring the amount of dye consumed by flies exposed to food hydrated with water or a copper solution within a 24-hour period. Both DSPR strain and treatment significantly influenced feeding (DSPR Strain: F_94,384_ = 3.08, P < 0.00001; Treatment: F_1,384_ = 2306, P < 0.00001, Table S2G; Figure 2A), although treatment had a much larger effect on the feeding response than strain (Cohen’s F: Treatment = 2.57, DSPR strain = 0.91). We also observed an interaction between strain and treatment (DSPR Strain x Treatment: F_94,384_ = 2.77, P < 0.00001, Cohen’s F = 0.86, Table S2G), indicating that the reduction in feeding due to copper is not uniform across strains (Figure 2). Both feeding responses had high heritability (control feeding response: H^2^ = 87.6%; copper feeding response: H^2^ = 87.6%).

**Figure 2.**
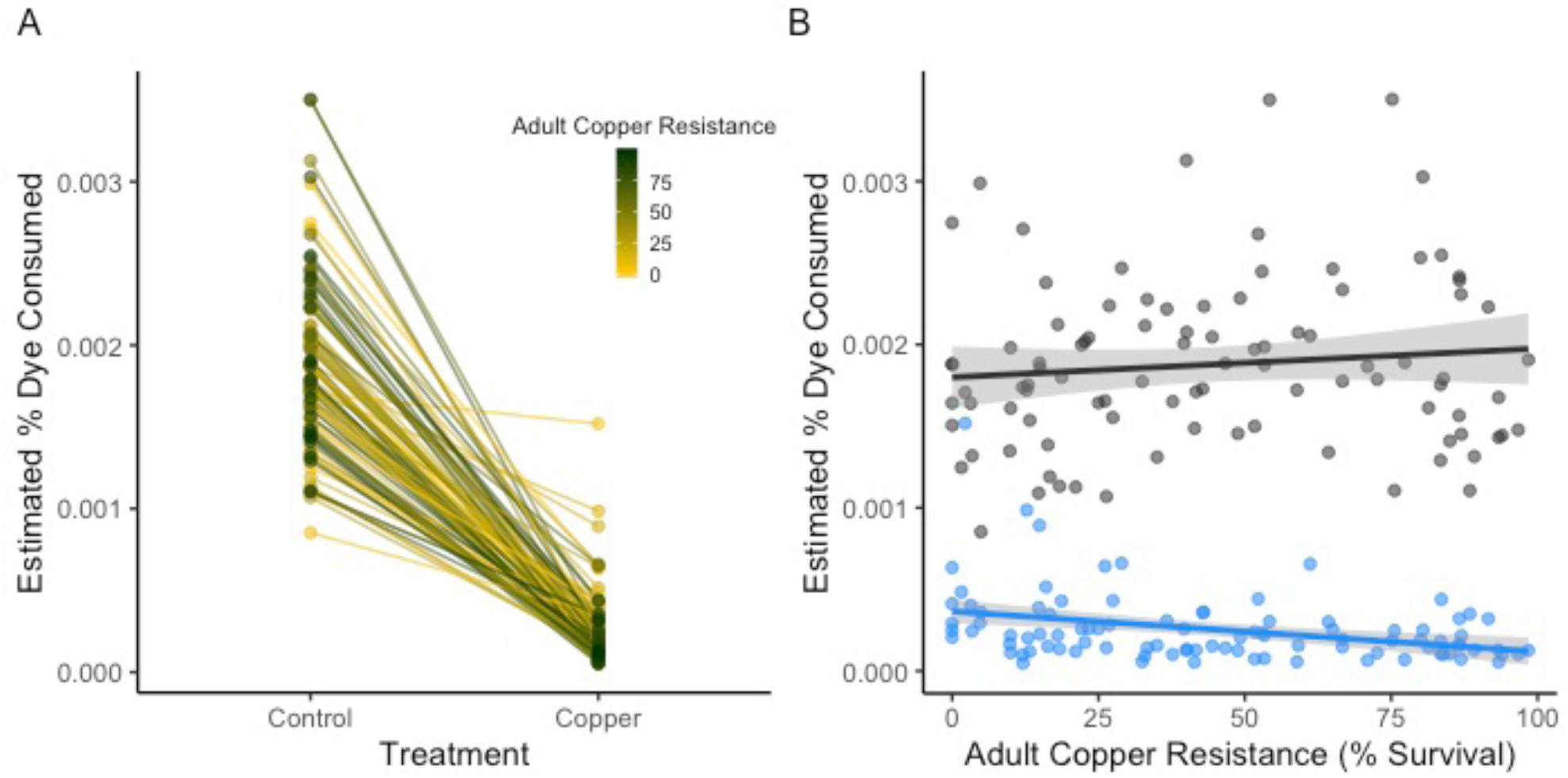
Feeding behavior in 95 DSPR strains changed in response to 24-hour exposure to 50mM CuSO4. A. Mean percent dye consumed varied among DSPR strains (P < 0.00001) and was much lower under copper conditions relative to control (water) conditions (Treatment: P < 0.00001). The interaction between strain and treatment (P < 0.00001) suggests that sensitivity to copper varies among the strains. B. Feeding behavior under control conditions was not correlated with adult copper resistance (P = 0.32); feeding behavior on copper was correlated with adult copper resistance (Adult Copper Resistance x Treatment: P = 0.03). Feeding response to copper is shown in blue; the control response is shown in black. Shading indicates the 95% CI of the regression.

Overall, feeding behavior under copper conditions was weakly negatively correlated with adult copper resistance (R = −34.6%, F_3,186_ = 263.7, P = 0.03; Table S2H, Figure 2B), while feeding behavior under control conditions was not correlated with adult copper resistance (R = 10.2%, F_1,93_ = 0.98, P = 0.32, Figure 2B). This suggests that our adult copper resistance phenotype is partially influenced by a copper-specific behavior, where more sensitive strains tend to eat more copper food than more resistant strains in a 24-hour period. Equally, the limited strength of the relationship likely implies our resistance phenotype is primarily impacted by the physiological and metabolic response to copper and is not solely influenced by behavioral avoidance.

### Copper-specific developmental response

In organisms with complex life cycles, the genetic control of physiological traits can be decoupled between life stages (Freda et al. 2017; Collet and Fellous 2019). To assess whether the strains with high resistance to copper as adults were also more resistant in other life stages, we sampled 100 strains from the B1 and B2 subpanels that span the range of adult copper resistance (from 0% – 98.4% survival). Embryos from these strains were placed on either control media, or media containing 2mM CuSO_4_, and the day of first adult emergence (development time) and the proportion of embryos that emerged as adults (developmental viability) was recorded. Both development time and developmental viability were variable among strains on copper and control media (development time: F_99,1157_ = 24.21, P < 0.00001; developmental viability: F_99,1157_ = 49.17.21, P < 0.00001; Table S2I,J, Figure 3). Exposure to copper delayed emergence by nearly 4 days on average (F_1,1157_ = 1293, P < 0.00001; Table S2I, Figure 3A) and significantly reduced developmental viability (F_1,1157_ = 3905, P < 0.00001; Table S2J, Figure 3B). There was an interaction between strain and treatment for both measures of the developmental response to copper, indicating that although development time and developmental viability were negatively affected by copper exposure for the majority of strains, the magnitude of the effect of treatment varied among strains (development time: F_1,1157_ = 11.75, P < 0.00001; developmental viability: F_1,1157_ = 13.13, P < 0.00001; Table S2I,J, Figure 3A,B)

**Figure 3.**
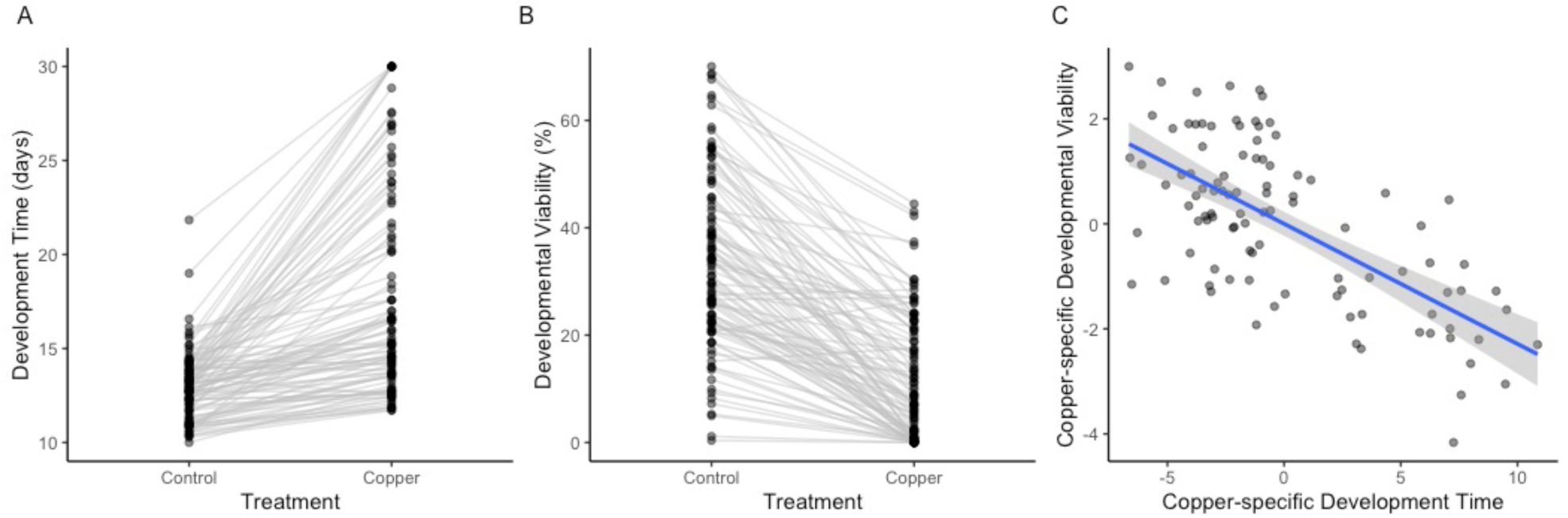
Development time (A) and developmental viability (B) were reduced in most strains by exposure to 2mM CuSO4. C. Copper-specific developmental viability and development time (corrected for strain-specific variation in these responses on control food) were correlated (P < 0.00001, R^2^ = 44%), indicating that strains with longer development time on copper-supplemented media also had lower viability. Points indicate the strain mean under each treatment condition. Grey shading indicates the 95% CI of the regression.

Because we were primarily interested in the effects of copper on development time and developmental viability, we regressed out variation under control conditions from both developmental phenotypes (see Materials and Methods). Copper-specific development time and copper-specific developmental viability were correlated (F_1,98_ = 76.4, P < 0.00001, R^2^ = 44%; Table S2K, Figure 3C), demonstrating that strains with longer development time on copper also had lower viability. Heritability was similar between copper-specific development time (H^2^ = 87.7%) and copper-specific developmental viability (H^2^ = 87.7%).

Neither copper-specific development time nor copper-specific developmental viability were correlated with adult copper resistance at an alpha level of 0.05 (copper-specific development time: F_1,98_ = 0.16, P = 0.69, R^2^ = 0.02; Table S2L, Figure 4A; copper-specific developmental viability: F_1,98_ = 2.71, P = 0.10, R^2^ = 2.7%, Table S2M, Figure 4B). The lack of a significant correlation between both measures of the developmental copper response and adult copper resistance could imply that copper resistance is influenced by life stage-specific mechanisms. However, because several other aspects of our assays used to measure the adult and developmental response to copper differ (e.g. copper concentration and exposure time), the lack of a significant correlation between these life stage specific measures of the response to copper stress is likely also influenced by technical variation.

**Figure 4.**
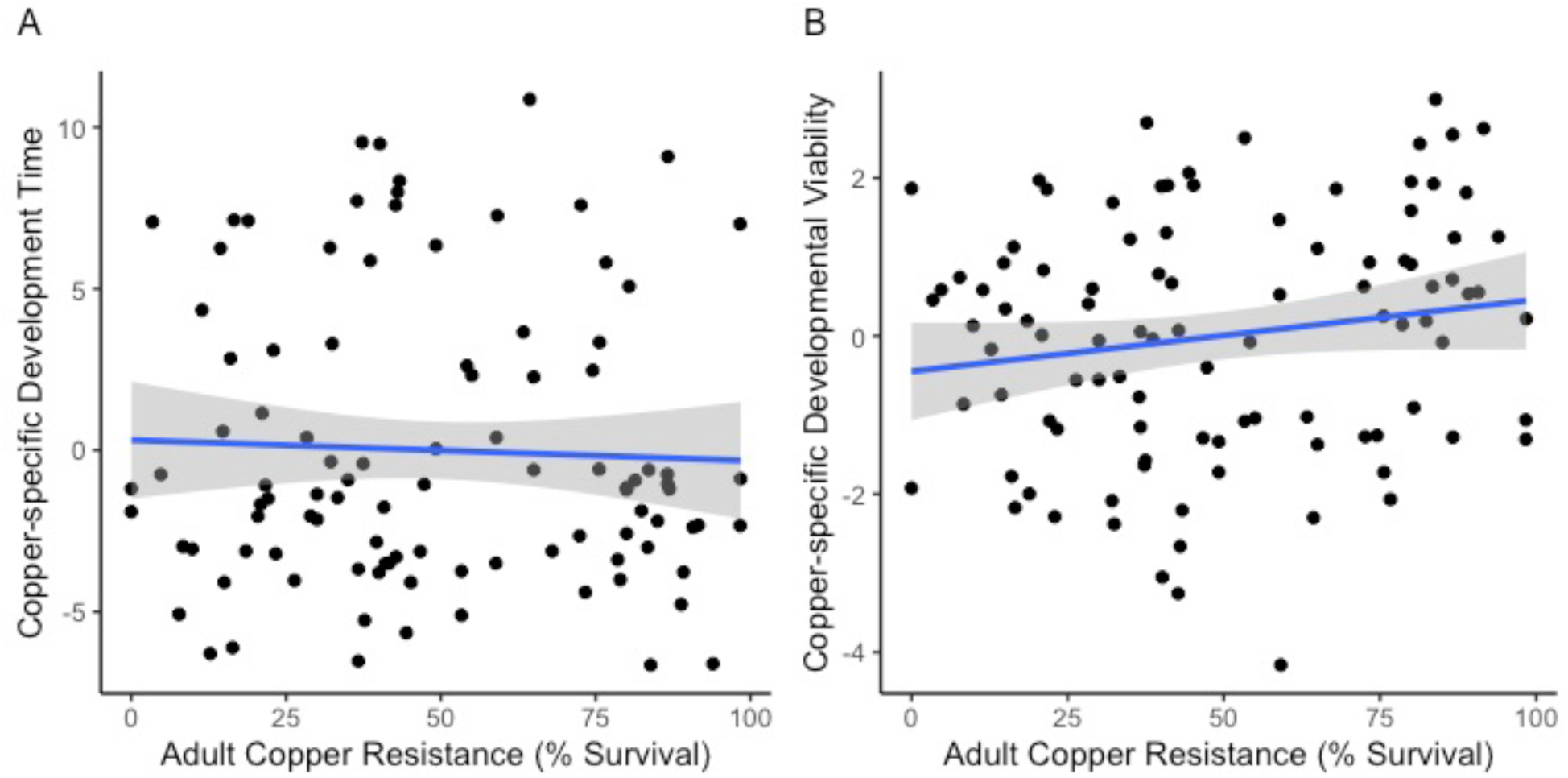
Copper resistance is not correlated across life stages. A. Copper-specific development time was not correlated with adult copper resistance (P = 0.69). B. Copper-specific developmental viability and adult copper resistance were not correlated (P = 0.10). In both plots, points indicate strain means. Higher, positive values for copper-specific development time and developmental viability indicate longer or higher development time or viability on copper, respectively. Grey shading indicates the 95% CI of the regression between residual developmental response (corrected for variation in the response on control food) and adult female survival after 48hrs on 50mM CuSO4.

### QTL mapping of life stage-specific response to copper stress

A principal goal of our study was to genetically dissect the response to copper stress, and using the DSPR we identified a total of 12 QTL between the A and B panels that were associated with variation in the adult copper resistance (Figure 5A, Table 1, Figure S6). Assuming that each QTL contributes to phenotypic variation in an additive manner, the QTL explained a substantial amount of variation in adult copper resistance (A panel: 36.19%; B panel: 27.93%; Table 1). The genetic architecture of adult copper resistance was largely panel-specific, with only one QTL (Q3) overlapping between the mapping panels. The 2-LOD drop interval of Q3 in the A panel fell entirely within the 2-LOD drop interval of Q3 in the B panel (Table 1, Figure 5A). Panel-specific genetic architecture of trait variation is consistent with several other studies that have mapped traits in both panels of the DSPR (Marriage et al. 2014; Najarro et al. 2017; Everman et al. 2019).

**Figure 5.**
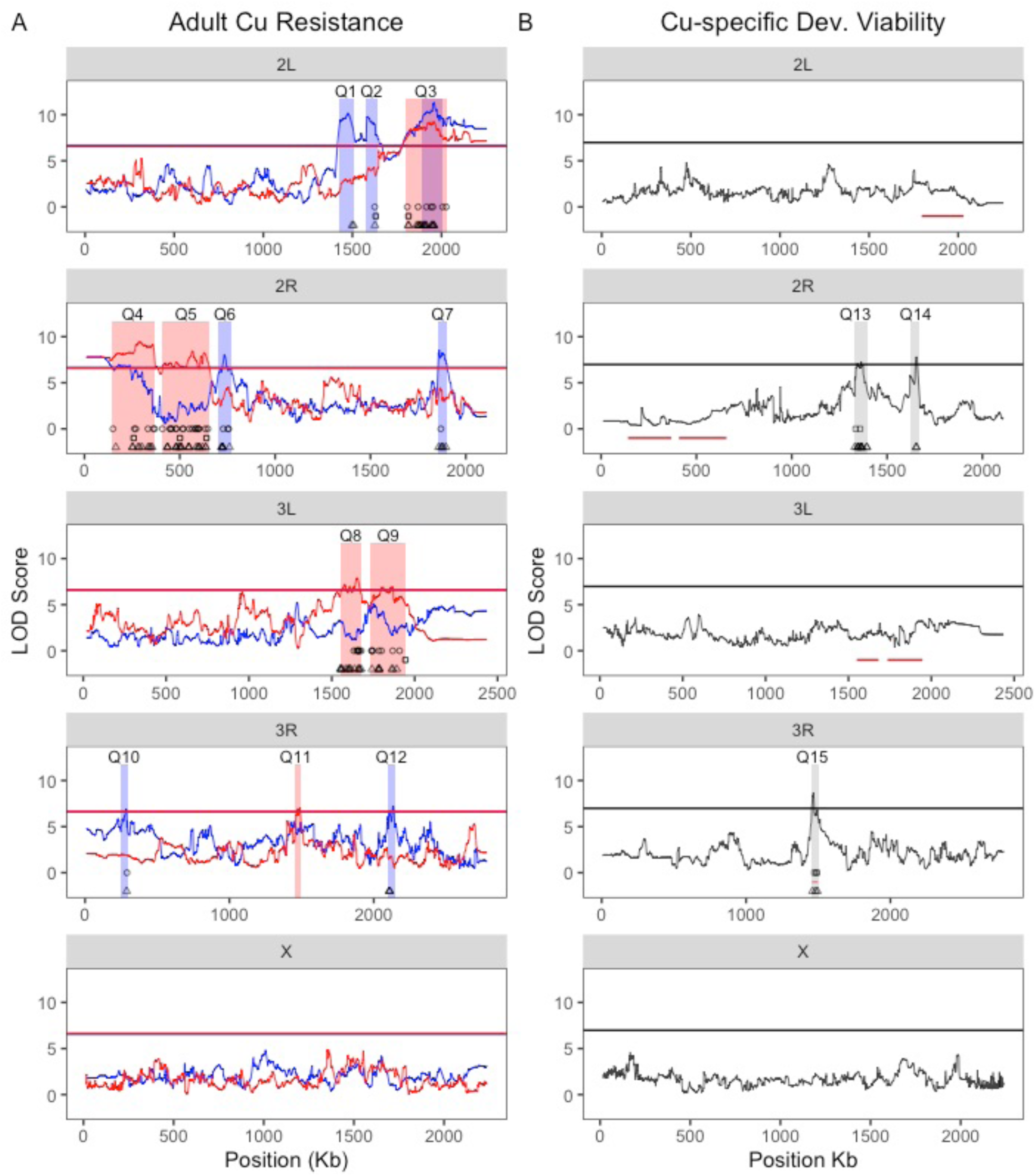
QTL associated with variation in adult copper resistance (A) and copper-specific developmental viability (B). A. We detected several QTL in the A (blue) and B (red) DSPR panels. Most QTL were panel specific with one QTL (Q3) overlapping between panels. Red and blue bars represent the 2-LOD drop intervals for each QTL. B. QTL mapped for copper-specific developmental viability. One QTL (Q15) for developmental viability overlapped with Q11 contributing to adult copper resistance. Red horizontal lines represent the 2-LOD drop intervals for the 6 QTL associated with the B panel adult survival response to copper, and grey bars represent the 3-LOD drop for the three QTL associated with copper-specific developmental viability (*Table 1*). The horizontal lines in each plot represent permutation-derived 5% critical thresholds (the thresholds for each panel in A. are nearly identical, leading to the lines overlapping.) Round points indicate DE genes influenced by resistance class, triangle points indicate DE genes influenced by treatment, and square points indicated DE genes that are shared between the treatment and resistance class models.

This lack of QTL peak overlap is likely the result of using a different set of founders to establish each mapping panel (King and Long 2017) but may also reflect a lack of power (King et al. 2012a) or epistatic effects that influence our ability to detect all QTL underlying adult copper resistance in each panel.

We did not find any QTL contributing to food consumption on either control or copper-supplemented food or copper-specific development time using both a strict (*α* = 0.05) and relaxed (*α* = 0.2) significance threshold, but we did find three QTL that contributed to variation in copper-specific developmental viability (Figure 5B). Given we only phenotyped 100 DSPR strains, power deficits certainly contribute to the low numbers of QTL identified for these traits (King et al. 2012a). Additionally, due to Beavis effects (Beavis et al. 1991; King and Long 2017), estimates of QTL effects based on 100 DSPR strains are typically overestimated (King and Long 2017), so the relatively high estimates of the variance explained by the QTL mapped for developmental viability (Table 1) should be interpreted with care.

One copper-specific developmental viability QTL (Q15) overlapped with a QTL (Q11) associated with adult copper resistance in the B panel (Figure 5). The 2-LOD drop interval of Q11 fell entirely within the 3-LOD drop interval of Q15. Multiparental mapping panels allow estimation of the effects of the founder haplotypes at mapped QTL. To assess whether the founder haplotypes contributed to adult copper resistance and copper-specific developmental viability in similar ways, we tested the correlation between the estimated founder effects at the shared QTL. Given that the location of the copper-specific developmental viability Q15 peak may be poorly estimated due to the low sample size employed for mapping, we compared the haplotype effects from the adult and developmental datasets at the peak position of the Q11 adult copper resistance QTL. These estimated founder effects were significantly positively correlated (F_1,5_ = 7.11, P = 0.04, R^2^ = 59%; Table S2N, Figure 6), suggesting that the alleles at the estimated QTL peak position of Q15 and Q11 influence the response to copper stress in adults and developing individuals in a similar way. This result implies that the adult and developmental response to copper stress are not fully independent, as was suggested by the very weak phenotypic correlation between these traits (Figure 4).

**Figure 6.**
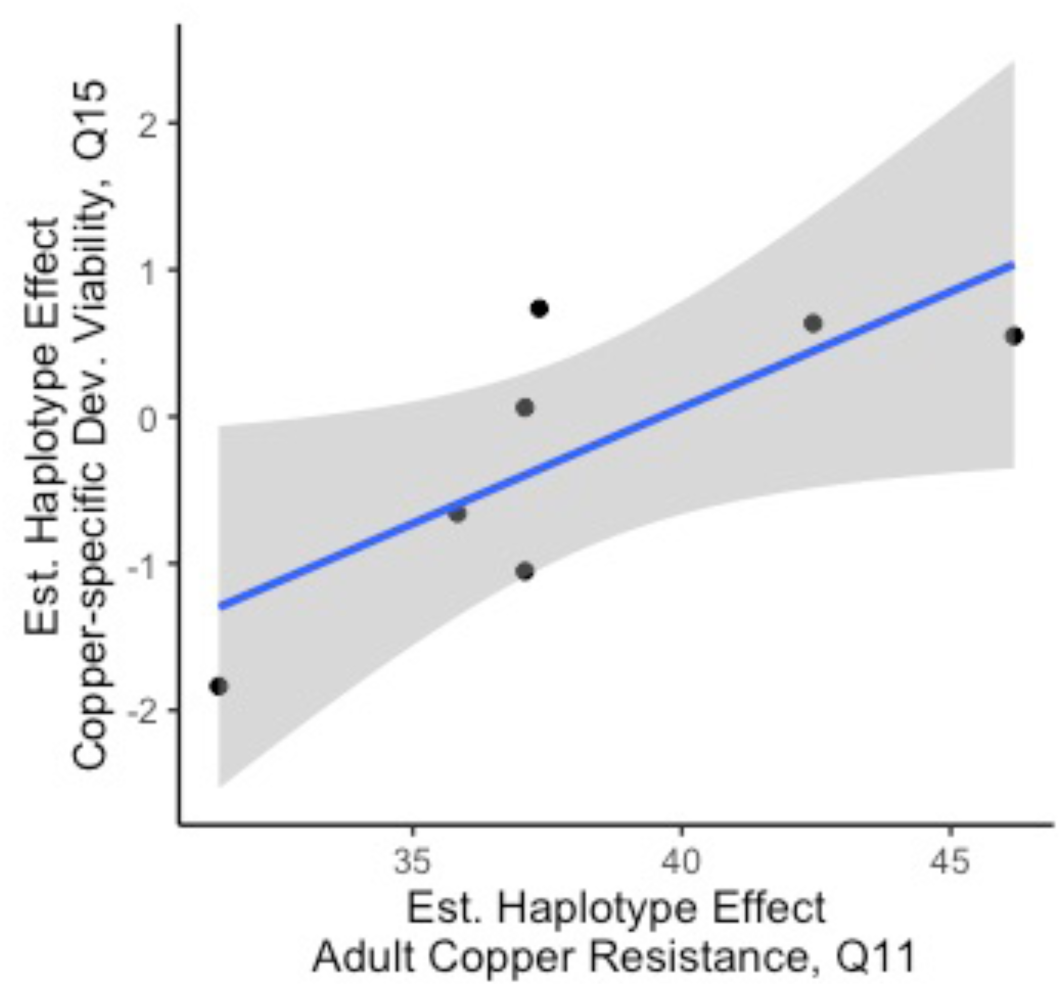
Founder haplotype effects for copper-specific developmental viability and adult copper resistance estimated at a shared QTL position on chromosome 3R were significantly positively correlated (P = 0.04, R^2^ = 59%). Grey shading indicates the 95% CI of the regression between estimated founder haplotype effects at Q11 for adult copper resistance and at the equivalent genomic position for copper-specific developmental viability, which resides within Q15.

### Genes implicated by mapped QTL

Combined across panels, the QTL regions associated with adult copper resistance include a total of 1823 unique protein coding genes. Of these, 10 genes have been previously associated with copper homeostasis, binding, chaperone activity, or copper cell development (Table S4). Promising copper-associated candidate genes include *Syx5*, *Grx1*, *CG11825*, *Ccs*, *Sod3*, and *CG5235*. *Syntaxin 5* (*Syx5*), associated with Q2, *Glutaredoxin 1* (*Grx1*), associated with Q9, and *CG11825*, associated with Q5, are all thought to be play a role in copper ion homeostasis (Norgate et al. 2007, 2010; Mercer and Burke 2016). *Syx5* is required for normal uptake of cellular copper, and it plays a critical role in copper ion homeostasis in *D. melanogaster* that is independent of other copper transporter proteins such as *Ctr1A/B* (Norgate et al. 2010). Similarly, *Grx1* knockdown results in copper deficiency, and this gene may function as a mediator of copper transfer to chaperone proteins (Mercer and Burke 2016). *CG11825* has been identified as a candidate for copper ion homeostasis in *D. melanogaster* by (Norgate et al. 2007), but functional testing is lacking for this gene under copper stress conditions. The gene copper chaperone for superoxide dismutase (*Ccs*), found under Q5, is an important chaperone protein that shuttles copper ions to *Sod1* under normal conditions (Culotta et al. 1997; Schmidt et al. 2000). Genetic ablation of *Ccs* in *D. melanogaster* resulted in increased sensitivity to oxidative stress following paraquat exposure (Kirby et al. 2008); however, the effect of *Ccs* knockdown under copper stress conditions has not been assessed. Genes previously associated with copper ion binding include *Sod3* (Q6) and *CG5235* (Q8). While *Sod3* functions as an extracellular receptor for copper ions and is protective against oxidative stress (Blackney et al. 2014), the link between *CG5235* and copper is based only on prediction informed by gene ontology (Gaudet et al. 2011). In addition to these copper-associated genes, we observed 64 genes with functions related to homeostasis or detoxification of zinc, 2 genes involved with manganese regulation, and 19 genes involved in binding unspecified metals. Of particular interest among these genes are *Catsup* (Q3), *ZnT41F* (Q4), and *stl* (Q7), which are all associated with zinc transport or detoxification (Yepiskoposyan et al. 2006; Ozdowski et al. 2009; Lye et al. 2013; Navarro and Schneuwly 2017), *trpl* (Q5) and *DCP2* (Q8), which are hypothesized to be involved in manganese ion binding (Thurmond *et al*. 2019), and *swm* (Q3), *babo* (Q5), and *whd* (Q5), which are thought to be involved in binding of unspecified metal ions based on gene ontology prediction (Gaudet *et al*. 2011; Thurmond *et al*. 2019).

The three QTL associated with copper-specific developmental viability spanned a total of 247 unique protein coding genes. Of these genes, none had functions previously linked to copper. However, 8 genes were associated with zinc ion binding, and two were linked to metal ion binding through gene ontology prediction (Table S4; (Gaudet *et al*. 2011; Thurmond *et al*. 2019)). Most notable among the genes identified by copper-specific developmental viability QTL was *mekk1* (Q15), which was demonstrated though gene knockdown to be the primary activator of JNK signaling under cadmium stress in *Drosophila* S2 cells (Ryabinina et al. 2006). Although the Q15 developmental viability QTL overlaps with the adult copper resistance QTL Q11, *mekk1* is only present within the interval implicated by Q15. Given that *mekk1* is within 11.2Kb of the Q11 2-LOD drop interval, this gene may still be a plausible candidate for adult copper resistance.

We performed GO enrichment analysis with genes implicated by adult copper resistance QTL in both the A and B panels as well as with genes implicated by QTL associated with copper-specific developmental viability with FlyMine (Lyne et al. 2007). No GO enrichment was observed for genes included in QTL intervals for either panel or life stage. This is unsurprising given that QTL intervals include many genes that are likely to be non-causative and potentially obscure any signal of enrichment.

### Differential gene expression due to treatment and resistance class

Allelic effects on variation in complex traits are commonly mediated by regulatory variation (Roelofs et al. 2006; Ruden et al. 2009; Boyle et al. 2017; GTEx Consortium et al. 2018). We used an RNA sequencing approach to examine the effects of copper stress on gene regulation and to assess any differences in this response between genotypes with high or low adult copper resistance. We sequenced mRNA from whole females from 6 high (79 – 98%) and 4 low (0 – 18%) adult copper resistance strains from the B panel following a 9-hour exposure to control (water) and 50mM CuSO_4_ conditions. A primary goal was to determine whether there are consistent differences in gene expression between high and low resistance classes of strains when exposed to copper stress, so we treated each strain as a replicate of the high and low resistance classes.

The interaction between treatment (control vs 50mM CuSO_4_) and resistance class was not significant at a 5% FDR or at a relaxed cutoff of 20%, so this term was dropped from the model and the treatment (TRT) and resistance class (RES) terms were assessed additively. After additional filtering (see methods), we identified 1589 genes that were differentially expressed across treatment and adult copper resistance class with the full model (full model: ∼ TRT + RES vs reduced model: ∼ 1). We used PCA with quantile-normalized filtered TPM data from these 1589 genes to explore the patterns of gene expression among the sampled strains and treatments, finding a pronounced effect of gene expression on treatment, with a more subtle effect on resistance class (Figure 7, Table S5).

**Figure 7.**
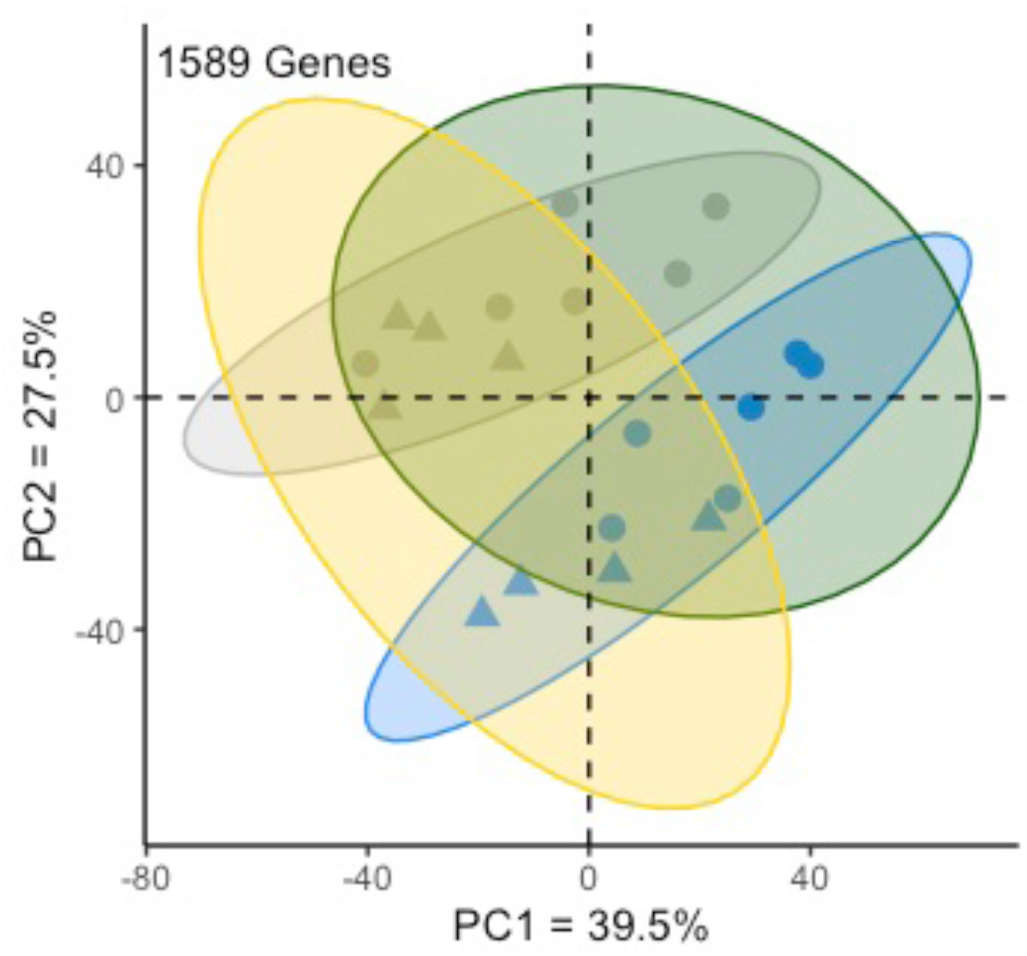
Principal components analysis of significantly differentially expressed genes (quantile-normalized filtered TPM) identified by the full model (full model: ∼ TRT + RES vs reduced model: ∼ 1). The effect of treatment was pronounced among samples, while the effect of resistance level was more subtle. Ellipses indicate the equivalent of a 95% confidence interval. Blue indicates copper-exposed samples, grey indicates control-exposed samples, green indicates resistant strains, yellow indicates sensitive strains. Triangle points indicate sensitive strains; circles indicate resistant strains.

To further explore how each of the main effects of the differential expression model influence gene expression, and to identify sets of genes that were influenced primarily by treatment or resistance class, we tested the effects of treatment and resistance class separately. Treatment alone (treatment model: ∼ TRT + RES vs. reduced model: ∼ RES) primarily contributed to differential expression in 848 genes, and adult copper resistance alone (resistance model: ∼ TRT + RES vs. reduced model: ∼ TRT) primarily contributed to differential expression in 466 genes. The vast majority of genes influenced by treatment and resistance class were included among the 1589 genes identified with the full model (92% of genes identified with the treatment model, 91% of genes identified with the resistance model). Of the 848 and 466 genes identified with the treatment and resistance class models, 58 genes were shared. The proportion of shared genes increased when a more relaxed significance threshold (20% FDR) was used in the treatment and resistance class models, and the estimated effects of resistance class and treatment on gene expression in each DE gene list were weakly positively correlated (Treatment Model: R^2^ = 8%; Resistance Class Model: R^2^ = 2%). While the DE gene lists attributed to treatment and resistance class are not fully independent, they represent sets of genes for which the primary source of variation is either treatment or resistance class.

To broadly characterize DE genes identified in the treatment and resistance class models, we performed GO analysis with FlyMine (Lyne et al. 2007) for each gene list separately. GO analysis of each complete DE gene list is summarized in Table S6. Briefly, 111 GO terms were identified from the full model DE gene list and included terms related to cell organization, cell cycle, and metabolism among the top 10 (Table S6). Enrichment for 23 GO terms including those related to cytoplasmic translation, ribosome biogenesis, and RNA processing was observed for the 848 genes influenced by treatment (Table S6). Fifty-nine GO terms were identified from the 466 DE genes due to resistance class. Top among these GO terms were those related to ATP synthesis, cellular respiration, and mitochondrial function (Table S6). GO analysis of this set of 58 genes revealed enrichment for female gamete generation [*GO:0007292*] (P = 0.006), and no genes had any connection to copper or metal ion homeostasis to our knowledge. Many of the genes with DE due to treatment and/or resistance class fell within the QTL intervals for adult copper resistance or copper-specific developmental viability (Figure 5). Notably, no enrichment was observed for any GO term related to metal ion homeostasis or detoxification when the total DE gene lists were considered (but see below). Of the 848 genes with DE due to treatment, 87 (11%) overlapped with QTL intervals, and of the 466 genes with DE due to resistance class, 62 genes (13.3%) overlapped with QTL intervals. Of the 58 genes shared between the treatment and resistance class models, 12 genes (20.6%) overlapped with QTL intervals (Figure 5). DE genes from the treatment and resistance class models were not more likely than expected by chance to fall within QTL intervals (*χ*^2^ = 14.5, df = 14, P = 0.41), although this does not preclude the possibility that those DE genes within mapped QTL are strong candidates to carry variation contributing to copper resistance.

To further explore the influence of resistance class on gene expression, we calculated the average change in gene expression following copper exposure for each of the 1589 DE genes from the full model using the same filtered TPM data used for PCA for the high and low resistance classes. The absolute values of these data were then log transformed to reduce spread, and the sign of the change in gene expression was restored by multiplying the result by 1 or −1. Copper-induced genes had higher expression under copper conditions, while copper-repressed genes had lower expression under copper conditions.

Of the 848 DE genes identified in the treatment model, there was a roughly even split between induced and repressed genes, with no difference between the resistance classes (Kolmogorov-Smirnov (KS) test: D = 0.05, P = 0.33; Figure 8A). Among the top 20 most highly induced genes under copper conditions in both resistant and sensitive classes were several MTs (*MtnA*, *MtnC*, *MtnD*, *MtnE*) as well as two genes that comprise a major iron storage complex (*Fer1HCH* and *Fer2LCH*). Because these genes and other genes with DE due to treatment were induced in sensitive and resistant strains to similar degree, we suggest that sensitivity to copper is not due to a failure to induce expression of genes with protective functions against copper ions. Among the 466 DE genes identified in the resistance model, gene induction by copper was more frequently observed in sensitive strains compared to resistant strains (KS test: D = 0.27, P < 0.00001; Figure8B). The top 20 most highly induced genes under copper conditions in sensitive strains included several genes that are involved in mitochondrial structure, function, and energy synthesis (e.g. *Ald1*, *levy*, *sesB*, *Mpcp1*, *COX5A*, *ATPsynb*), suggesting more sensitive strains may be characterized by a greater susceptibility to oxidative stress.

**Figure 8.**
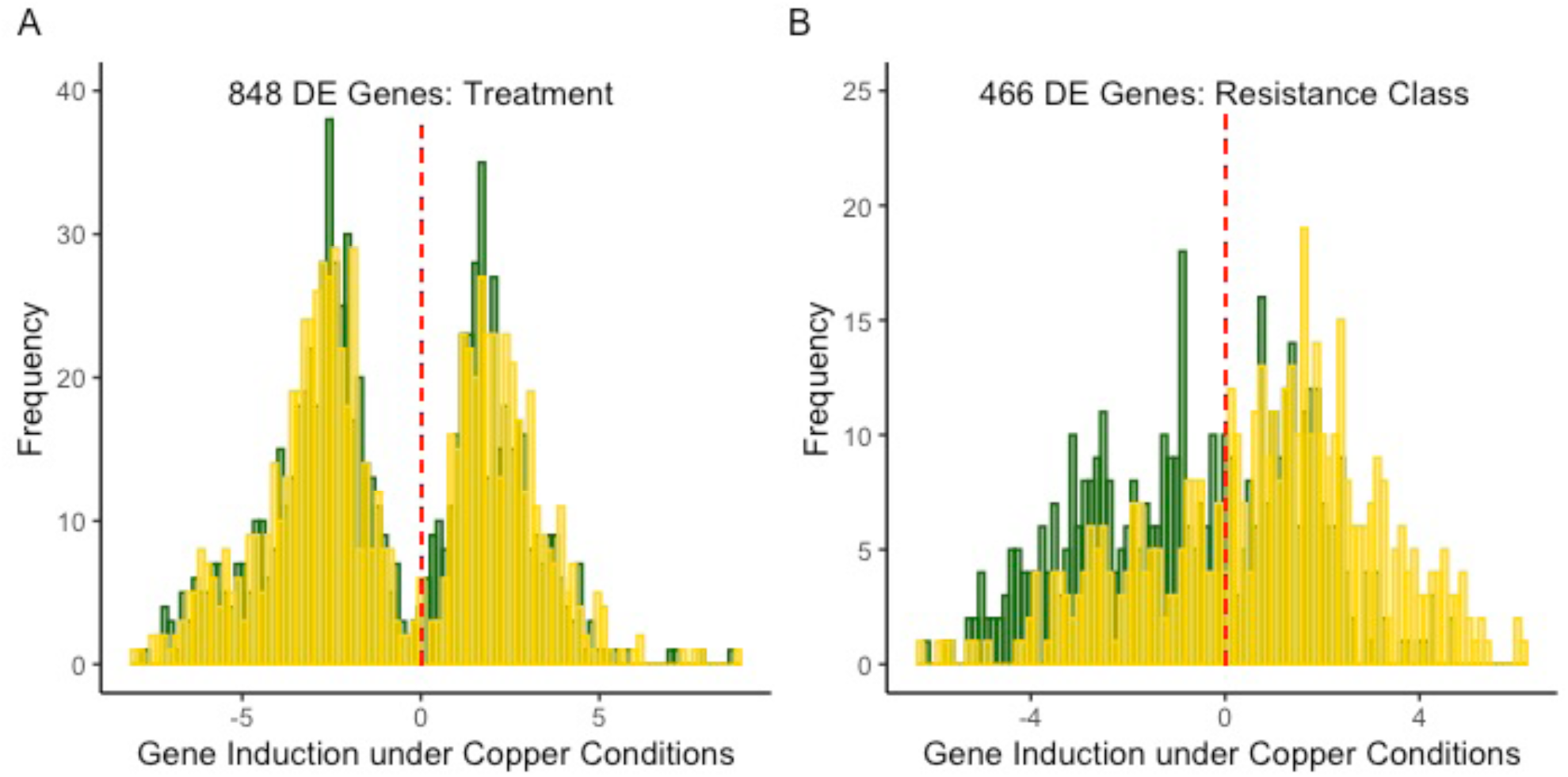
Induction of genes under copper conditions with DE identified from the treatment model (A) andf the resistance model (B). A. The effect of treatment on DE highlights that roughly equal numbers of genes were induced or repressed under copper conditions (KS-test: D = 0.05, P = 0.33). B. Among DE genes identified by the resistance model, genes were more likely to be induced by copper exposure in sensitive strains compared to resistant strains (KS-test: D = 0.27, P << 0.001). In each plot, yellow bars indicate gene expression in sensitive strains and green bars indicate gene expression in resistant strains.

### Cluster analysis of differentially expressed genes

The patterns of copper-induced gene expression across treatment and resistance class observed in Figure 8A and B raise the question of whether there are co-regulated sets of genes that distinguish resistant and sensitive strains under copper and control conditions. For example, the presence of metal-associated genes among the those induced by treatment may signal a larger network of genes that are co-regulated in response to heavy metal stress. To identify any such co-regulated groups, we used Clust (Abu-Jamous and Kelly 2018) to identify non-overlapping clusters of genes from the 848 genes influenced by treatment and the 466 genes influenced by resistance class using quantile-normalized, filtered TPM data. We then performed GO analysis on any resulting clusters formed from the two sets of genes with FlyMine (Lyne et al. 2007).

Clust identified 3 clusters of co-regulated genes with DE due to treatment (Figure S7A). Treatment clusters 1 (101 genes) and 2 (17 genes) consisted primarily of genes that were induced by copper exposure, while treatment cluster 3 (17 genes) consisted solely of genes that were copper-repressed (Figure S7A). While top GO terms for treatment clusters 1 and 3 revealed enrichment for genes involved in processes unrelated to metal ion homeostasis or response (e.g. cell cycle, RNA processing; Table S6), treatment cluster 2 was enriched for genes involved in iron import, transport, and detoxification of iron and inorganic compounds (Table S6). Gene enrichment in treatment cluster 2 is consistent with our expectation that copper-induced genes will include those that are involved in the response to toxic metal ion exposure. Interestingly, only one of the genes (Gclc) in treatment cluster 2 has been previously directly associated with copper through its interactions with the copper transport proteins *Ctr1A* and *ATP7*. Also included in treatment cluster 2 are *Fer1HCH* and *Fer2LCH*, which are primarily involved with iron storage but may interact with copper ions during protein assembly (Huard et al. 2013). Treatment clusters 1 and 3 included 17 and four genes, respectively, that were implicated by adult copper resistance-associated QTL (Figure S7A). The gene *dnk* in treatment cluster 1 was also implicated by the copper-specific developmental viability QTL Q15 (Figure S7A). Two of the genes (*CG11878* and *CG5506*) identified in treatment cluster 2 were implicated by adult copper resistance-associated QTL intervals (Figure S7). One gene, *CG5506*, was empirically demonstrated to interact with *Fer2HCH* (Guruharsha et al. 2011); however, neither gene has been previously associated with copper exposure.

Clust identified 2 clusters of co-regulated genes that were differentially expressed due to resistance class (Figure S7B). Resistance cluster 1 (75 genes) was primarily enriched for genes involved in cell cycle processes (Table S6), and 11 genes were also implicated by adult copper resistance QTL. Resistance cluster 2 (56 genes) included genes that were more often copper-induced in sensitive strains and copper-repressed in resistant strains (Figure S7B). Resistance cluster 2 was enriched for two broader categories of GO terms including several related to muscle structure (e.g. myofibril assembly [*GO:0030239*], P < 0.00001) and mitochondrial function and energy synthesis (e.g. ATP metabolic process [*GO:0046034*], P < 0.00001). Resistance cluster 2 also included genes involved in inorganic ion homeostasis [*GO:0098771*], although enrichment for this GO term was weak (P = 0.05). Of the genes involved in inorganic ion homeostasis, three (*CG14757*, *trpl*, and *sesB*) are particularly noteworthy given that all three genes are copper-induced in sensitive strains and copper-repressed in resistant strains, and two have been previously linked to metal ion homeostasis (Figure 9). Exposure of the Drosophila S2 cell line to 2mM CuSO_4_ resulted in increased expression of *CG14757*, indicating that this gene is responsive to copper stress (Norgate et al. 2007); however, its exact function relative to the toxic effects of copper has not been elucidated. The gene *trpl* is predicted to be involved in manganese ion binding (Thurmond *et al*. 2019) and was included in the adult copper resistance-associated QTL Q5 in this study. *sesB* is a mitochondrial transporter gene that was demonstrated to be important for protection against oxidative stress through gene knockdown in D. melanogaster (Terhzaz et al. 2010). Other genes included in this group (*up*, *SERCA*, and *nrv3*; Figure 9) are involved in transport of calcium, sodium, and potassium (Domingo et al. 1998; Gaudet et al. 2011) or are thought to be involved in ATP metabolism (*Vha68-1*) (Thurmond *et al*. 2019). In addition to *trpl*, 2 other genes from resistance cluster 2 were implicated by adult copper resistance QTL.

**Figure 9.**
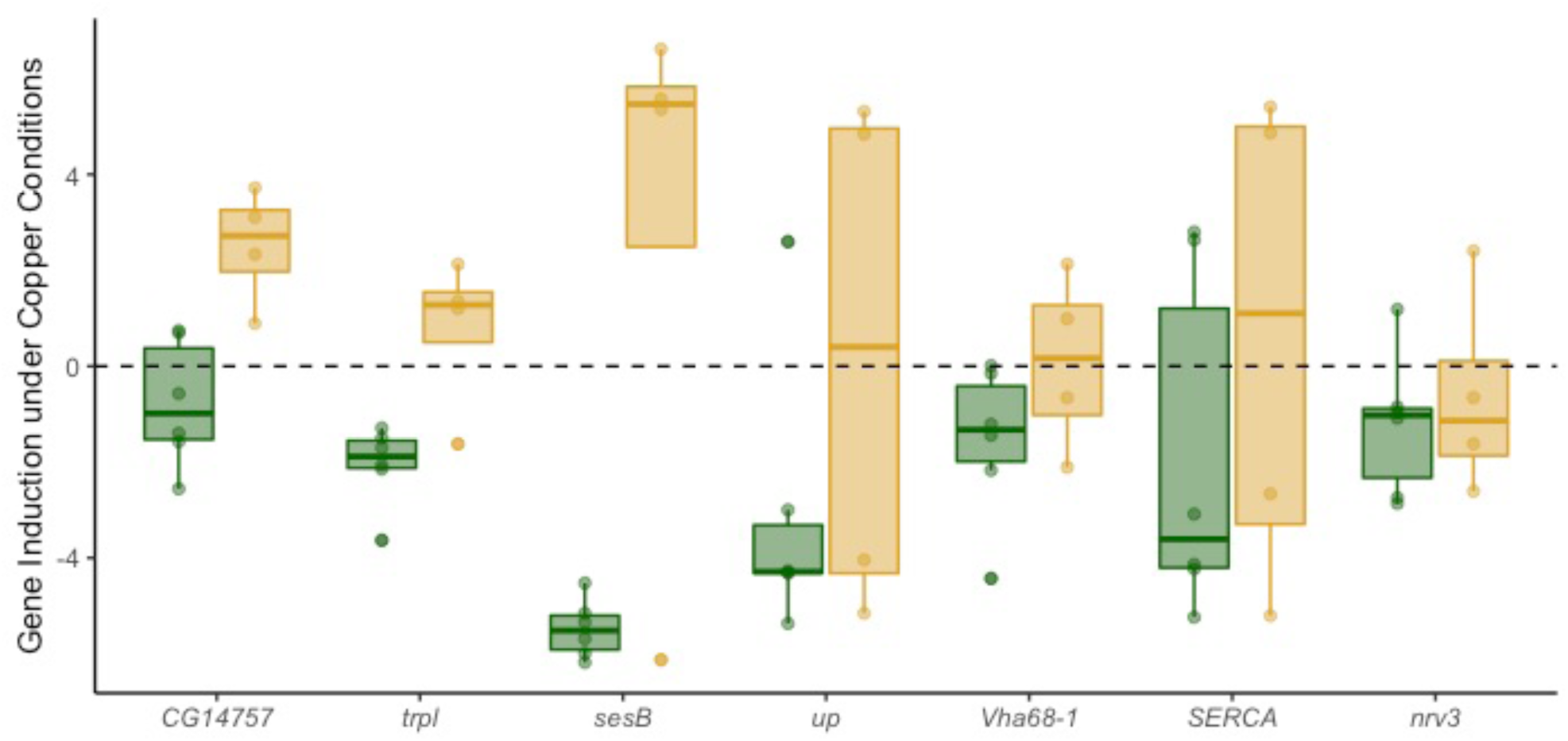
Copper-induced expression of genes involved in inorganic ion homeostasis that were included in resistance cluster 2. Resistant strains are shown in green, sensitive strains are shown in yellow.

### RNAi knockdown of candidate genes associated with adult copper survival

Several genes with links to copper or metal ion homeostasis were implicated by QTL or were differentially expressed due to treatment or resistance class (see above) (Table S3). We picked 16 genes to functionally test using RNAi knockdown. QTL-implicated genes included *Catsup* and *swm* (Q3), *ZnT41F* (Q4), *CG11825*, *whd*, *babo*, and *Ccs* (Q5), *stl* (Q7), *DCP2* and *CG5235* (Q8). We also tested *trpl* (Q5), which, along with *Mvl*, was among the DE genes influenced by resistance class. Because *Ccs* and both *Sod1* and *Sod2* closely interact, we tested *Sod1/2* even though these genes were not implicated by either QTL or RNAseq. From genes with DE due to treatment, we tested *MtnC* and *CG10505*. Of these candidate genes, only *Sod1*, *MtnC*, and *Mvl* have been previously specifically linked to copper stress (Calap-Quintana et al. 2017). *Ccs*, *CG5235*, *Sod1*, *CG11825*, and *CG10505* are all associated with copper transport or binding. The remaining candidate genes (*trpl*, *DCP2*, *whd*, *stl*, *swm*, *babo*, *Catsup*, and *ZnT41F*) have not been experimentally linked to copper stress, but are associated with metal ion binding or homeostasis.

Genes were tested using TRiP UAS RNAi strains (Perkins et al. 2015) that were crossed to a background with a ubiquitously expressed Gal4-expressing driver resulting in knockdown in the whole animal and to a background with an anterior midgut-specific Gal4-expressing driver resulting in knockdown in this specific region of the midgut. We measured average lifespan of all knockdown genotypes on 50mM CuSO_4_ with at least 10 replicates.

In general, more genes influenced copper resistance when they were knocked down in the whole animal compared to when candidates were knocked down in the anterior midgut (Figure 10). Of the candidate genes with known associations with copper, *Ccs*, *CG5235 (b)*, *MtnC (b)*, and *Sod1* reduced copper resistance relative to the control when knocked down in the whole animal using the ubiquitous driver (Figure 10A). Inconsistent effects of ubiquitous *CG5235* knockdown may be influenced by vector efficiency; the knockdown vector for *CG5235 (a)* is a long dsRNA vector (VALIUM10), while the knockdown vector for *CG5235 (b)* is a shRNA vector (VALIUM20). Both TRiP strains for *MtnC* used the same vector (VALIUM20) (Perkins et al. 2015); however, these two strains target *MtnC* at different locations within the gene, and knockdown efficiency may differ between the two sites. Knockdown of copper-associated genes in the anterior midgut did not influence copper resistance relative to the control, suggesting that reduced expression of *Ccs*, *CG5235*, *Sod1*, *CG11825*, and *CG10505* in this limited region of the midgut does not hinder the fly’s ability to cope with copper stress (Figure 10B).

**Figure 10.**
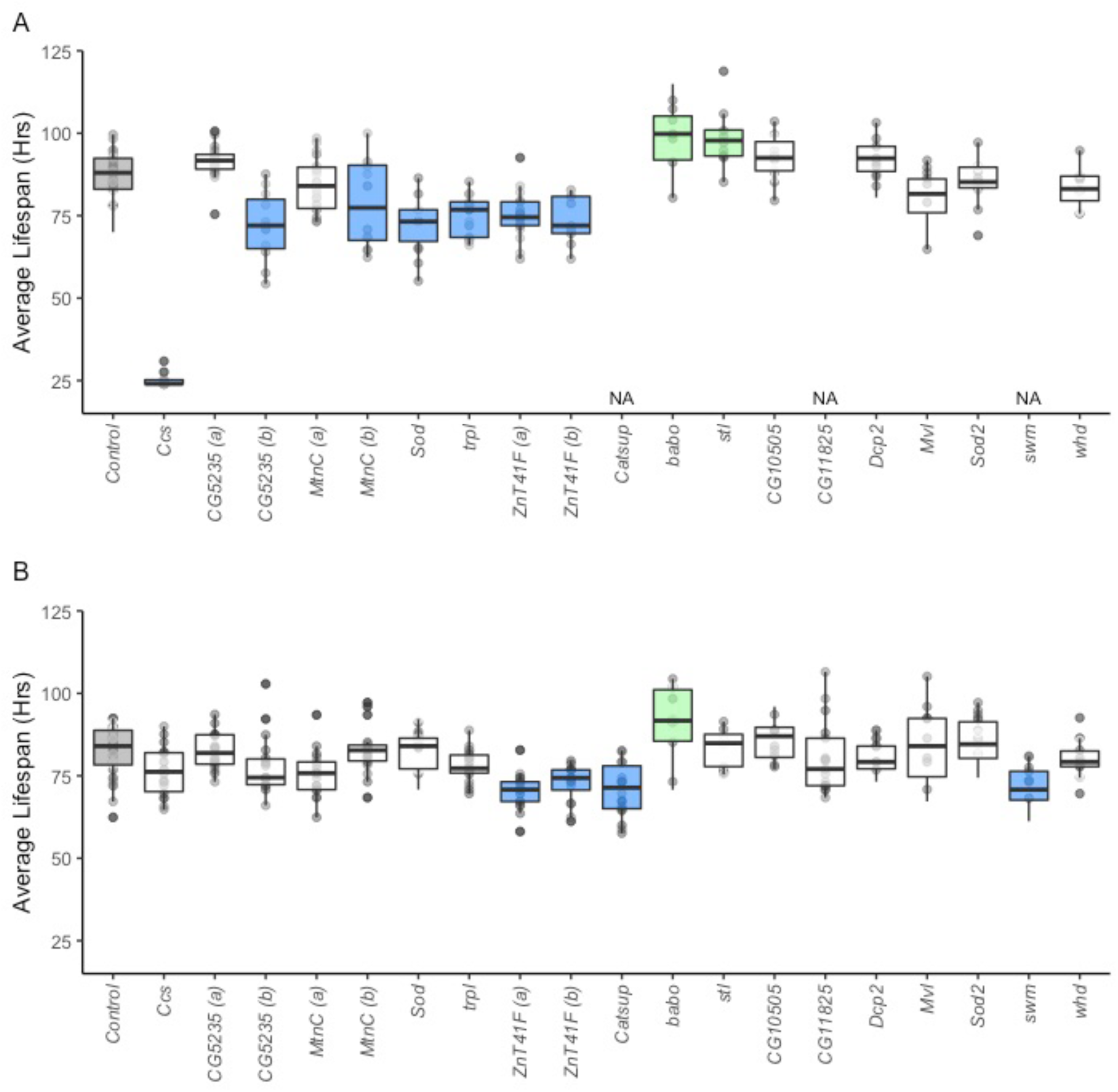
Average lifespan of TRiP UAS RNAi knockdown strains crossed to a ubiquitous Gal4-expressing driver (A) and to an anterior-specific Gal4-expressing driver (B). A. Increased susceptibility was observed with knockdown of Ccs, CG5235 (b), MtnC (b), Sod, trpl, and ZnT41F with the ubiquitous Gal4 driver. Knockdown of babo resulted in increased resistance to copper toxicity relative to the control. B. Knockdown in the anterior midgut of Catsup, swm, and ZnT41F resulted in increased susceptibility to copper toxicity, while knock down of babo increased resistance relative to the control. In each plot, grey shading indicates the control, green shading indicates increased resistance to copper, blue shading indicates decreased resistance, and no shading indicates lack of a significant difference based on an experiment-wide a = 0.05. Three candidate gene TRiP strains (swm, Catsup, and CG11825) produced too few flies to test when crossed to the ubiquitous Gal4-expressing driver, and were thus excluded from our analysis. We tested multiple TRiP UAS RNAi strains for genes CG5235 (CG5235 (a), CG5235 (b)), MtnC (MtnC (a), MtnC (b)), and ZnT41F (ZnT41F (a), ZnT41F (b)) to assess the consistency in the effect of gene knockdown on copper survival (*Table S3*).

The candidate gene *ZnT41F* consistently reduced copper resistance relative to control when knocked down in the whole animal and in the anterior midgut. While *ZnT41F* was previously shown to indirectly affect zinc homeostasis (Yin et al. 2017), the role it plays in copper ion homeostasis has not been described. Similarly, *Catsup* and *swm*, which have not been previously linked to copper, reduced copper resistance when knocked down in the anterior midgut. That knockdown of these genes in the whole animal did not influence copper resistance suggests these genes interact with copper soon after ingestion, although this would require additional follow-up to confirm. Interestingly, knockdown of *babo* in both the whole animal and the anterior midgut increased copper resistance relative to the controls (Figure 10). Knockdown of *stl* in the whole animal had a similar effect. Both genes are predicted to be involved in metal ion binding (Gaudet *et al*. 2011; Thurmond *et al*. 2019), but any additional evidence linking them to the detoxification of heavy metal ions under stressful conditions is lacking.

## Discussion

### Variation in heavy metal stress is influenced by a complex genetic architecture

*D. melanogaster* has been a particularly important model for elucidating the roles of genes involved in the response to copper and other heavy metals (e.g. Egli et al. 2003, 2006a; b; Balamurugan et al. 2004, 2007; Yepiskoposyan et al. 2006; Calap-Quintana et al. 2017). In our study, we used this model to investigate the role of genetic diversity in resistance to the heavy metal copper. We used a combination of QTL mapping and RNA sequencing to characterize allelic and gene expression variation that influences resistance to copper stress in strains from the multiparental DSPR mapping panel. In comparison with previous reports investigating the genetic architecture of copper and resistance to other heavy metals in plants (Macnair 1993; Selby and Willis 2018), the genetic architecture of copper resistance in *D. melanogaster* appears to be more complex. Where one to three QTL were identified for heavy metal resistance in several plant species including *Mimulus guttatus*, wheat, and corn (Allen 1971; Macnair 1983, 1993; MacNair et al. 1993; Bálint et al. 2007; Selby and Willis 2018), we identified 12 QTL that underlie variation in adult copper resistance (Figure 5A), and found that the diverse DSPR strains varied widely in survival following exposure to copper stress (Figure 1).

Part of the difference in apparent complexity underlying the response to copper is likely due to the higher power of our mapping panel, which employs a much larger number of genetically diverse strains coupled with higher genetic marker density compared to the mapping populations used in plant studies (e.g. Willems et al. 2007; and Courbot et al. 2007; King et al. 2012a). Secondly, the structure of the DSPR may be particularly conducive for detecting allelic variation in genes that influence the response to copper stress given the global sampling of founder strains used to generate the DSPR (King et al. 2012a), which may capture more of the natural variation for copper resistance than that present in any one natural population. Third, in contrast to natural populations which are often of interest because of their proximity to heavy metal pollution (e.g. Allen 1971; Macnair 1983; Ramirez et al. 2005; Turner et al. 2010; Wuana and Okieimen 2011; Wright et al. 2015; Arnold et al. 2016), the DSPR is naïve to any form of heavy metal selection or stress. Strong selection for heavy metal resistance could reduce variation at causative genes and lead to an apparent reduction in the complexity of resistance (Arnold et al. 2016).

The level of genetic complexity for copper resistance described in our study is consistent with reports of metal resistance in flies, yeast, and worms where measures of resistance were conducted in other heavy metal-naïve mapping populations. The DGRP (*Drosophila* Genetic Reference Panel (Mackay et al. 2012)), another large D. melanogaster mapping panel, was used to demonstrate a complex genetic architecture for heavy metal exposure through GWA (genome wide association) and extreme QTL mapping (i.e. sequencing and comparing pools of individuals with divergent phenotypes) in adult and developing life stages (Montgomery et al. 2014; Zhou et al. 2016, 2017). In these studies, tens to hundreds of genes have been implicated in natural genetic screens for lead (Zhou et al. 2016, 2017), cadmium (Zhou et al. 2017), and methylmercury (Montgomery et al. 2014). In *Saccharomyces cerevisiae* using several extreme QTL mapping pools, Ehrenreich et al. (2012) demonstrated that more than 20 distinct loci were associated with resistance to cadmium and nearly 40 loci were associated with copper resistance. In *Caenorhabditis elegans*, Evans et al. (2018) found 4, 6, and 6 QTL associated with the response to cadmium, copper, and silver, respectively.

Interestingly, each of the studies mentioned above generated largely distinct lists of genes associated with the responses to the heavy metals tested, and overlap between candidate genes from Zhou et al. (2016) and (2017), Montgomery et al. (2014), Ehrenreich et al. (2012), and Evans et al. (2018) and the current study is minimal as well (Table S4, Table S5). It is worth noting that none of these shared genes (Table S4, Table S5) have been previously associated with heavy metal toxicity or homeostasis and would require additional validation to determine whether they are causative for resistance to any heavy metals. However, overlap in candidate genes identified for the response to different heavy metals raises the possibility that a single gene or gene network may influence the response to multiple heavy metals. Consistency in genetic architectures underlying the response to chemical stressors has been examined using large *S. cerevisiae* and *C. elegans* mapping panels (Ehrenreich et al. 2010; Evans et al. 2018), and while there is some evidence for shared genetic architectures between traits, it is clear that each toxin response is influenced by a largely trait-specific genetic architecture. For example, out of 82 QTL that were identified for 16 different toxins ranging from heavy metals to cancer therapy drugs, Evans et al. (2018) found three QTL hotspots that were implicated for multiple types of toxins. Similarly, Ehrenreich et al. (2010) showed that ∼20% of QTL detected in their study were shared between 2 – 5 of 17 different chemical stressors in yeast. Only one QTL was shared between more than five different chemical stressors (Ehrenreich et al. 2010). In a later study Ehrenreich et al. (2012) demonstrated similarly low levels of consistency in genetic architecture through extreme QTL mapping of 13 chemicals. Although we did not measure resistance to multiple heavy metal stressors in our study, we functionally validated several candidate genes implicated by QTL and DE gene lists which had functions linked to other metals including zinc, lead, manganese, and cadmium, or that were not linked to a specific metal (e.g *swm* (Q3), *babo* (Q5), and *stl* (Q5); Figure 10). Pleiotropic gene effects inferred from our RNAi analyses may be the result of metal-sensitive genes responding to a generalized set of cytotoxic effects stemming from production of reactive oxygen species caused by heavy metal toxicity (Uriu-Adams and Keen 2005). This hypothesis is further supported by evidence of copper-induced expression of genes involved in oxidative stress response in sensitive strains that are repressed in resistant strains (e.g. *sesB*, Figure S7B, Figure 9). However, additional tests of the response of DSPR strains to a diverse set of heavy metals is needed to fully understand whether the non-copper candidate genes we identified have correlated effects on resistance to other heavy metals.

### Consistency in the genetic architecture of Copper resistance across life stages

Genes that are involved in copper homeostasis in *D. melanogaster* adults have in some cases also been shown to regulate copper in larvae. For example, exposure of larvae to CuSO_4_ induces expression of MTs (Egli et al. 2003, 2006b), and we demonstrated a copper-induced increase in MT expression in adults (Table S5). Knockdown of copper transporter genes in the CTR family alters copper homeostasis in both larvae and adult flies as well (Zhou et al. 2003; Turski and Thiele 2007). Given the similar genetic responses to copper in adults and larvae, we expected that the physiological response to copper stress would be correlated between life stages. It was previously established that developmental life stages are more susceptible to copper stress compared to adults; whereas adults were shown to survive for 80 days on 1mM CuSO_4_, developmental viability from egg to adult on 1mM CuSO_4_ was less than 10% in the same strain (Bahadorani and Hilliker 2009). Our goal was to understand the relationship between adult copper resistance and the effect of copper stress on development in a set of genetically diverse *D. melanogaster* strains.

Similar to previous reports, we found that copper stress delayed development and reduced viability (Zhou et al. 2003; Bahadorani and Hilliker 2009; Pölkki 2016) although to differing degrees among the DSPR strains (Figure 3), suggesting that as with adult copper resistance, copper-specific development time and developmental viability are genetically variable. Despite the lack of a statistically significant correlation between the developmental responses to copper stress and adult copper resistance (Figure 4), we did observe evidence of partially shared genetic architectures between copper-specific developmental viability and adult copper resistance (Figure 5B). Additional testing would be needed to determine whether the same genes implicated by developmental viability QTL Q15 and adult copper resistance QTL Q11 influence copper resistance at each life stage. Our detection of fewer QTL associated with copper-specific developmental viability may be directly related to the reduced number of genotypes sampled in our assay. Power to detect a 10% QTL with 100 DSPR strains is less than 20% (King et al. 2012a), so the actual level of overlap in the genetic architecture between the adult and developmental viability responses may be higher than we observe. Notably, founder haplotype frequencies in the full set of lines and the subset of 100 are very similar across the genome (Figure S8), so it is unlikely the case that the subset fails to capture the same allelic diversity present in the full set.

Because the ecology of the developing and adult stages of *D. melanogaster* are quite distinct, that copper resistance might be influenced by largely life stage-specific mechanisms is not unexpected. For example, *D. melanogaster* adults and larvae avoid copper-supplemented food when given the opportunity (Balamurugan et al. 2007; Bahadorani and Hilliker 2009); however, in natural populations, higher mobility of adults would allow the adult life stage to avoid heavy metal contaminated food more effectively. Life-stage specific genetic architectures were observed in *D. melanogaster* for cold tolerance (Freda et al. 2019), and the decoupling of the genetic mechanisms that influence survival and fitness have been reported in diverse organisms with complex life cycles (Moran 1994; Ragland and Kingsolver 2008). However, a number of other factors including the large difference in copper dose and the nature of the response tested at each life stage may obscure or complicate the relationship between the developmental and adult responses we observed. We used a much lower dose in our assessment of the effect of copper on development time and developmental viability compared to the adult copper resistance phenotype, and differences in dose can alter the genetic architecture for a trait. For example, in the yeast *S. cerevisiae*, Wang and Kruglyak (2014) demonstrated that the overall genetic architecture of haloperidol resistance was dose-dependent. While one QTL was consistently detected for each of the 5 doses tested, several QTL were only detected at a single dose (Wang and Kruglyak 2014). With the added complexity of assessing the effects of copper in different life stages, it is difficult to fully determine whether the effects of copper in the adult and developmental assays are analogous. Further confounding this comparison, our adult copper assay was implemented over a 48-hour time period in contrast to exposing developing flies from egg to adult to copper over a period of 30 days at most. The harmful effects of copper on development may be constant or variable across different stages (egg, larvae, pupation), and this represents an area of ongoing research.

### Copper sensitivity is influenced by gene expression variation and behavior

Differences in expression levels of genes that have protective functions against toxins, or that are co-opted by toxins can lead to variation in resistance levels. For instance in humans, natural variation in expression levels of the gene *CMG2* is associated with variation in resistance to anthrax (Martchenko et al. 2012), and in the fungus *Suillus luteus*, selection pressure from heavy metal pollution quickly led to copy number variation in transport genes with protective functions against heavy metal toxicity (Bazzicalupo et al. 2019). Sensitivity to copper in adult *D. melanogaster* DSPR strains does not appear to be due to insufficient expression of genes involved with copper or metal detoxification such as MTs or CTR family transporters (Figure S7, Table S5). Instead, we found that genes associated with metabolism and mitochondrial function were copper-induced in sensitive strains and copper-repressed in resistant strains (Figure S7B, Figure 9). Given that we also observed that sensitive strains are more slightly likely to consume copper in larger amounts in a 24-hour period compared to resistant strains (Figure 2), sensitive strains may be under greater metabolic stress as they cope with exposure to behaviorally-mediated higher levels of ingested copper. Copper resistance in *D. melanogaster* may not be simply a function of how well the organism is able to detoxify food; more likely, copper resistance is a combination of behavioral aversion to copper and the metabolic stress induced by the amount of metal consumed in addition to detoxification ability.

In general, food consumption rate has a complex genetic basis in *D. melanogaster* (Garlapow et al. 2015), and when given a choice, both *D. melanogaster* adults and larvae tend to avoid copper-supplemented food at much lower concentrations relative to those tested in this study (Balamurugan et al. 2007; Bahadorani and Hilliker 2009). Bahadorani and Hilliker (2009) showed that adult copper avoidance was observed at 1mM CuSO_4_, and avoidance in third-instar larvae was observed at 0.25mM CuSO_4_ (Balamurugan et al. 2007). Similarly, adult *D. melanogaster* avoid pupation and oviposition on copper-supplemented food (Bahadorani and Hilliker 2009). While this behavioral component likely plays an important role in mediating copper stress in natural populations, these studies focused on only one or few genetic strains, making it difficult to extrapolate how a genetically variable population would behave in response to copper. The correlation between adult copper resistance and copper food consumption in the 100 DSPR strains tested in our study suggests that variation in copper avoidance may play an important role in overall adult copper resistance. At this point, the specific relationship between copper consumption rates, metabolic stress, and genetic resistance to copper has not been characterized, but doing so in future studies has to potential to more clearly define resistance to ingested toxins compared to an assessment based solely on survival. Important remaining questions include whether behavioral avoidance and sensitivity to heavy metals are influenced by variation in chemosensory detection ability (e.g. Arya et al. 2015; He et al. 2016) or variation in preference for metal-supplemented food (e.g. Highfill et al. 2019). Addressing these questions with a large panel such as the DSPR will help support our efforts to characterize the relationship and potential interaction between behavior and genetic capacity for copper resistance.

## Conclusions

Copper resistance in *D. melanogaster* is genetically complex, is influenced by allelic and expression variation as well as by variation in behavioral avoidance of copper, and may be controlled by distinct sets of loci in different life stages. Several genes that have known copper-specific functions as well as genes that are involved in the regulation of other heavy metals were identified as potential candidates for variation in adult copper resistance and copper-specific developmental viability. We demonstrated that nine of these candidates influenced adult copper resistance, providing evidence of pleiotropic effects of genes previously thought to be associated with other heavy metals. Copper is just one of many heavy metals that pollute the environment with negative impacts on humans, fungi, plants, and insects at a global scale. Understanding the complexity of the genetic basis of copper resistance and the potential sources of variation that interact with resistance is important for understanding the diverse mechanisms through which copper pollution can negatively impact organisms.

## Acknowledgments

ERE was supported by Postdoctoral Fellowships from the NIH (F32 GM133111) and the Kansas INBRE project (P20 GM103418). The research was supported by NIH R01 OD010974 to SJM and Anthony Long (UC Irvine), and by NIH R01 ES029922 to SJM, and computational work was supported by the Kansas INBRE (P20 GM103418). Sequencing was carried out by the KU Genome Sequencing Core which is supported by the CMADP, an NIH COBRE award (P20 GM103638). Stocks obtained from the Bloomington Drosophila Stock Center (NIH P40 OD018537) were used in this study. We thank Robert Unckless and John Kelly for their comments on the manuscript, and William Jewell College for use of the macro camera lens used in the developmental assays.

## Supplemental Figures

**Figure S1.**
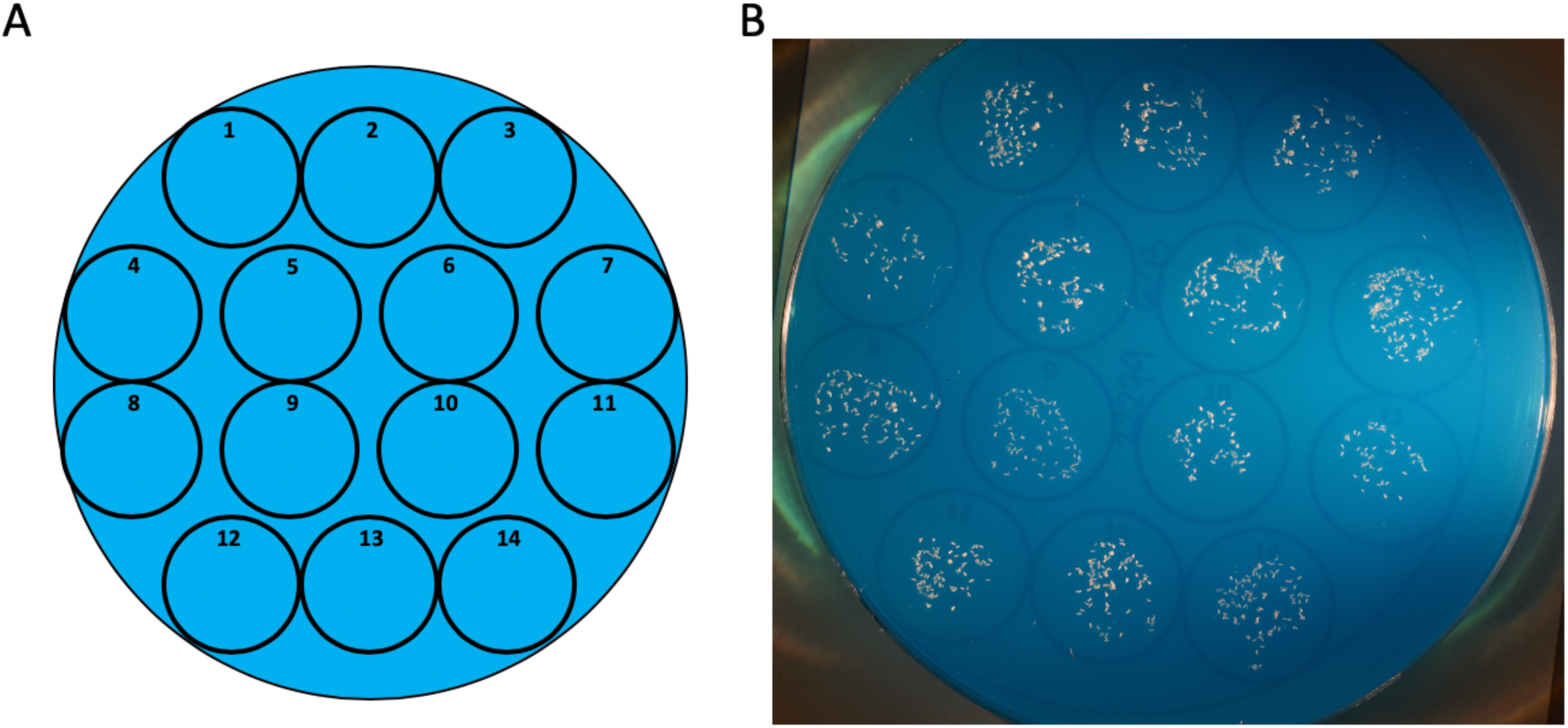
Embryos from each strain were arrayed on a 100mm petri dish containing 2% agar with dye for contrast. A. Template used for arraying embryos. B. An example of embryos arranged in each cell of the template. After taking a picture of each dish, embryos were transferred to vials containing water or 2mM CuSO4. All embryos from each cell on the template were transferred to one vial, resulting in up to 7 control vials and 7 copper vials per strain

**Figure S2.**
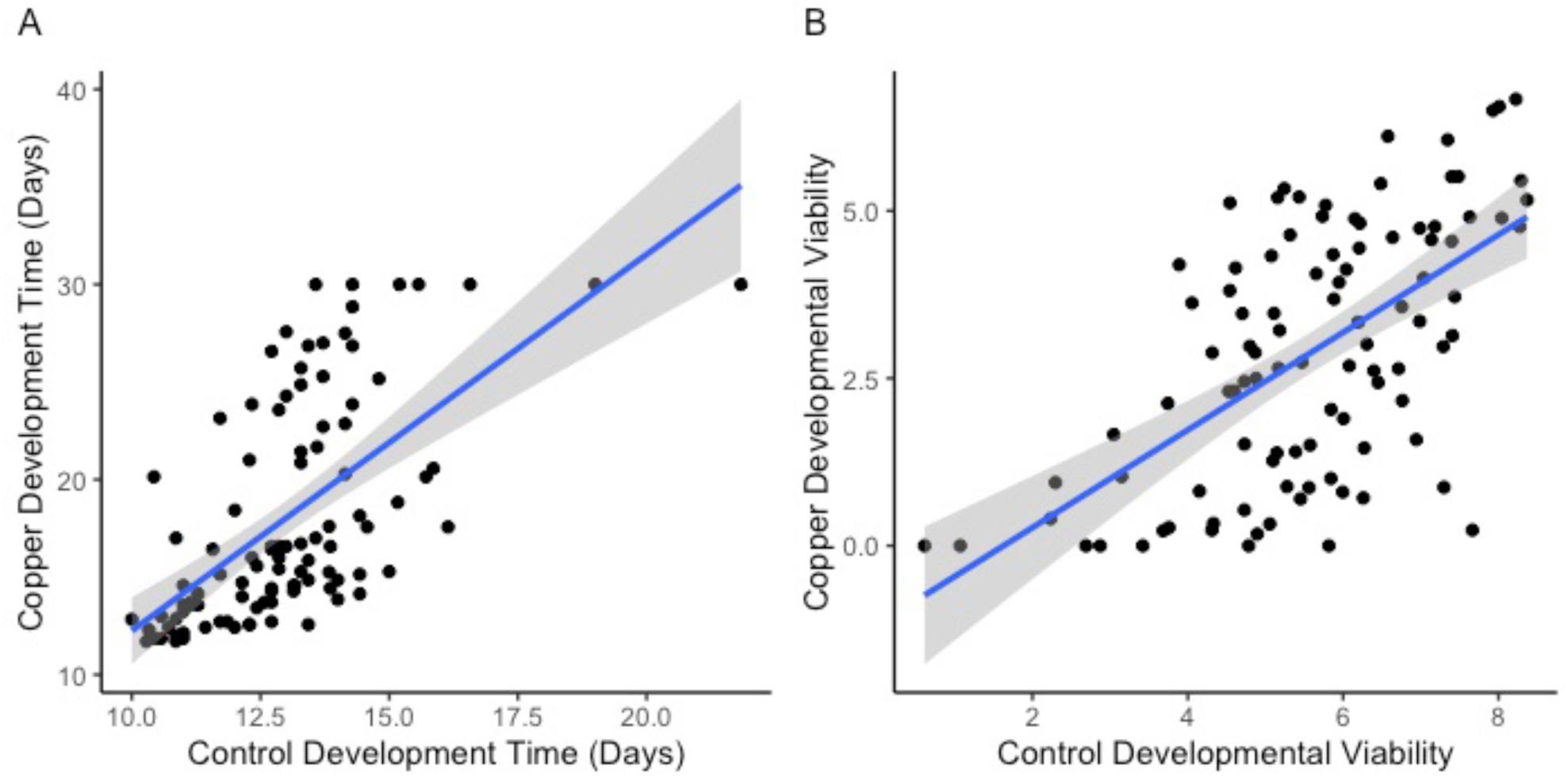
Correlation between developmental responses on control and copper media. A. Development time on control food was correlated development time on copper food (F(1,98) = 61.0, P < 0.0001, R^2^ = 38%). Vials in which no individuals emerged were given a value of 30 days. B Square root transformed developmental viability on control and 2mM CuSO4 was also correlated (F(1,98) = 57.1, P < 0.0001, R^2^ = 36%). Each point represents the strain mean response. Grey shading indicates the 95% CI of the regression.

**Figure S3.**
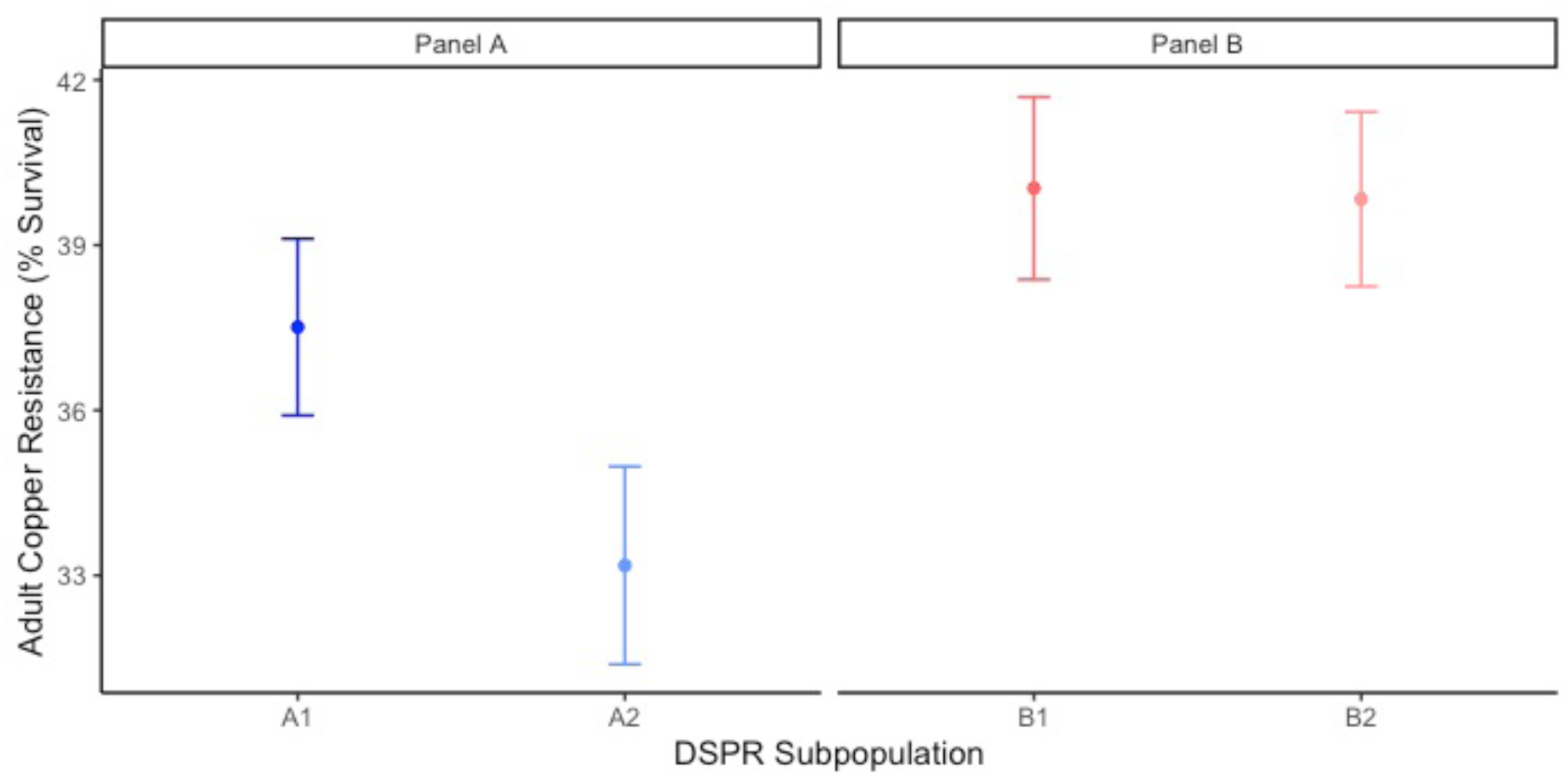
Mean (± 95% CI) female copper resistance per subpanel. Subpanel influenced percent survival in the pA (F1,2289 = 12.64; p < 0.001) but not pB panel (F1,2495 = 0.03; p = 0.86)

**Figure S4.**
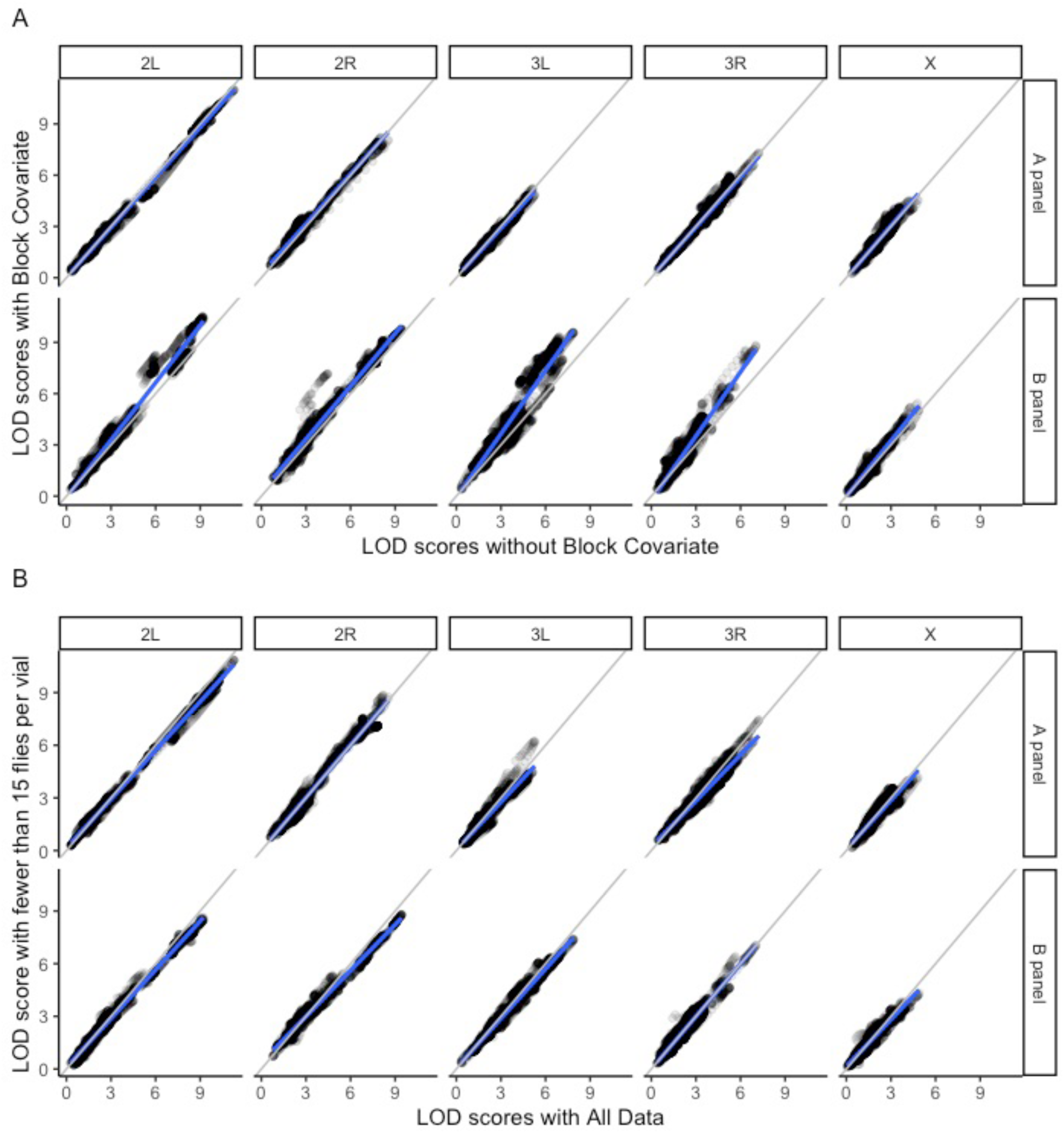
Comparison of LOD scores between a model with batch included as a covariate (A) and when vials with fewer than 15 flies were removed from the calculation of mean survival prior to mapping (B). Best fit lines are shown in blue in each plot; grey lines indicate the 1:1 line for comparison

**Figure S5.**
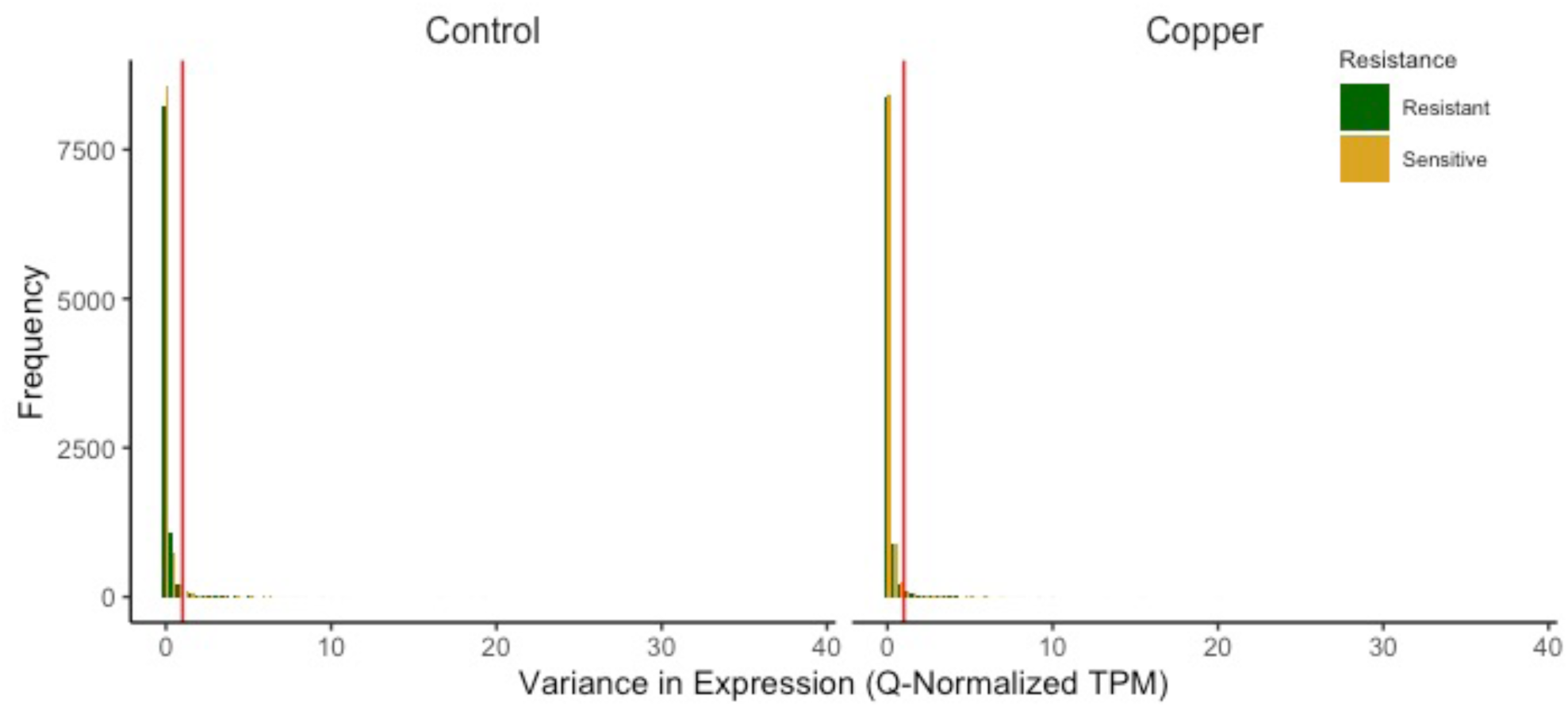
Variance in quantile-normalized TPM by within each sample group (resistant vs. sensitive) and treatment group (copper vs. control). The red vertical line indicates the cutoff of 1. All genes with a variance equal to or greater than 1 were excluded from all downstream analyses of DE gene from the three DE models

**Figure S6.**
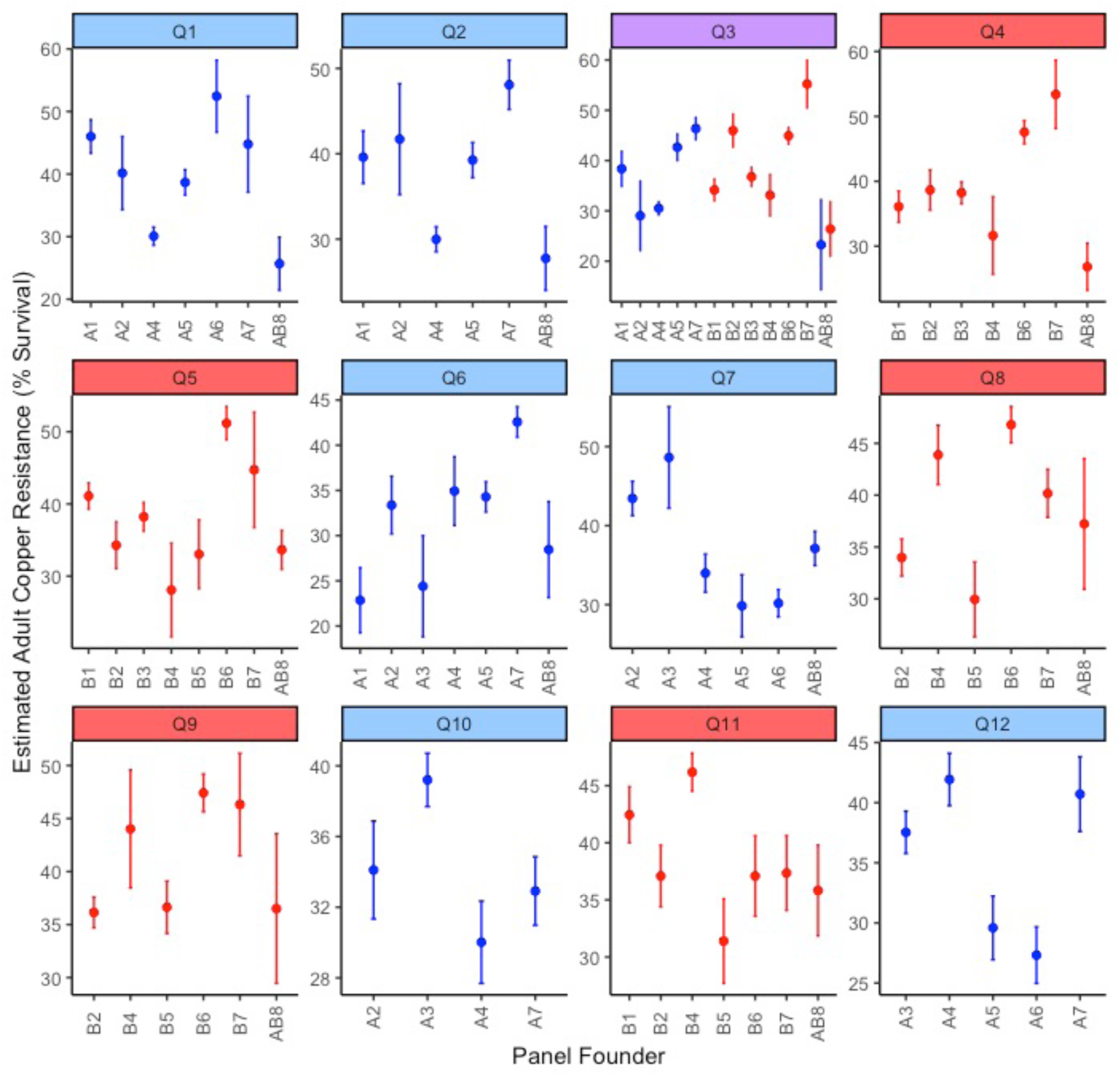
Estimated founder haplotype effects at each QTL associated with adult copper resistance. Data are presented as estimated founder means (±SE) when the founder haplotype was present in more than 7 DSPR strains. Plots are colored by panel; panel A is plotted in blue, panel B is plotted in blue. Purple indicates the shared QTL (Q3) between the A and B panels

**Figure S7.**
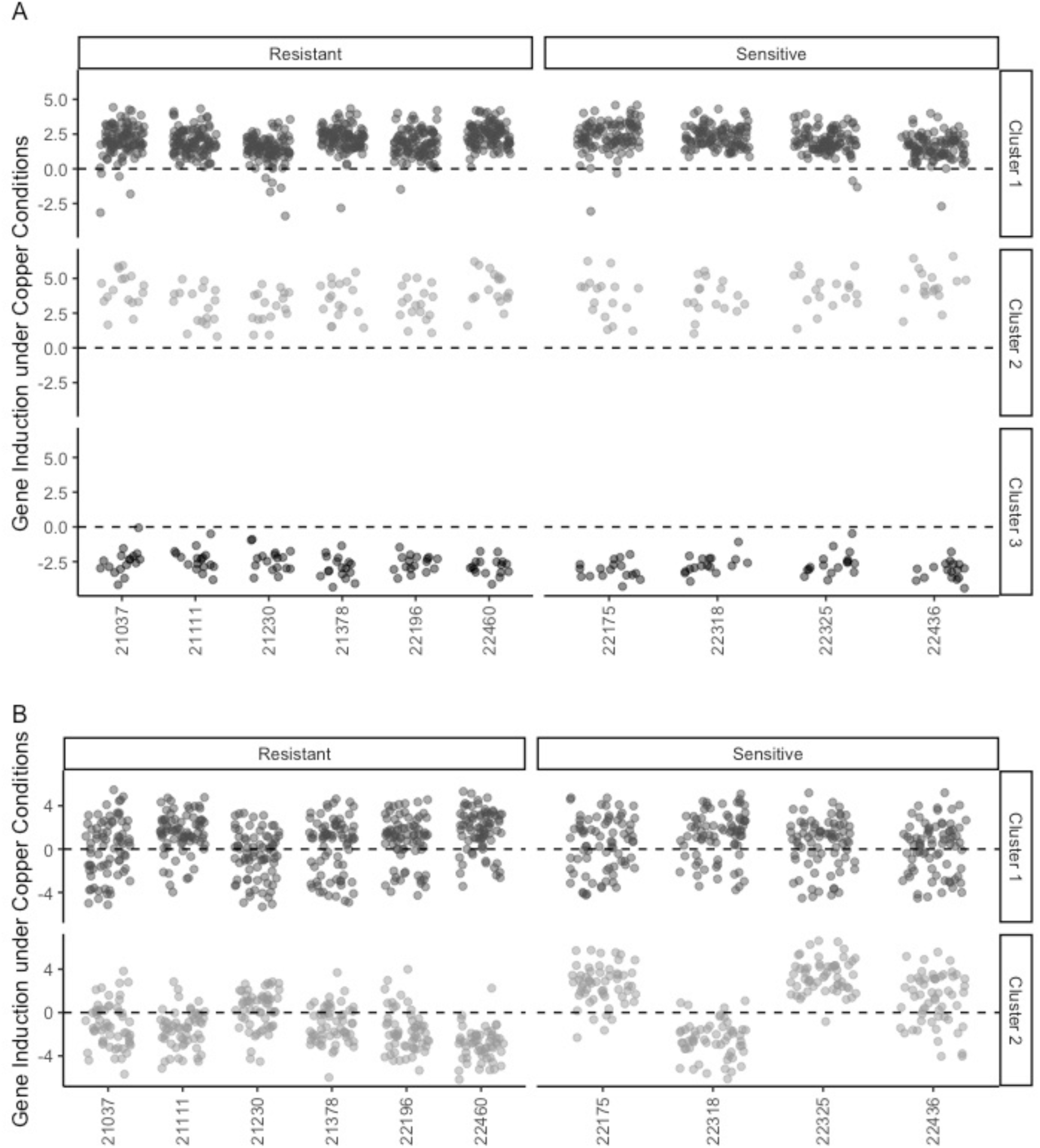
Clust analysis of genes influenced by treatment (A) or resistance class (B). A. Clust identified 3 clusters from genes with DE due to treatment. 17 genes from cluster 1 were also implicated by adult copper resistance QTL; one gene was also implicated by copper-specific developmental viability QTL Q15. From treatment clusters 2 and 3, 2 and 4 genes, respectively, were also implicated by adult copper resistance QTL. B. Clust identified 2 clusters from genes with DE due to resistance class. In cluster1, 11 genes were also implicated by adult copper resistance QTL; 4 genes from cluster 2 were also implicated by adult copper resistance QTL. One gene from cluster 2 was implicated by copper-specific developmental viability QTL Q15 as well. Points are shaded to help distinguish clusters

**Figure S8.**
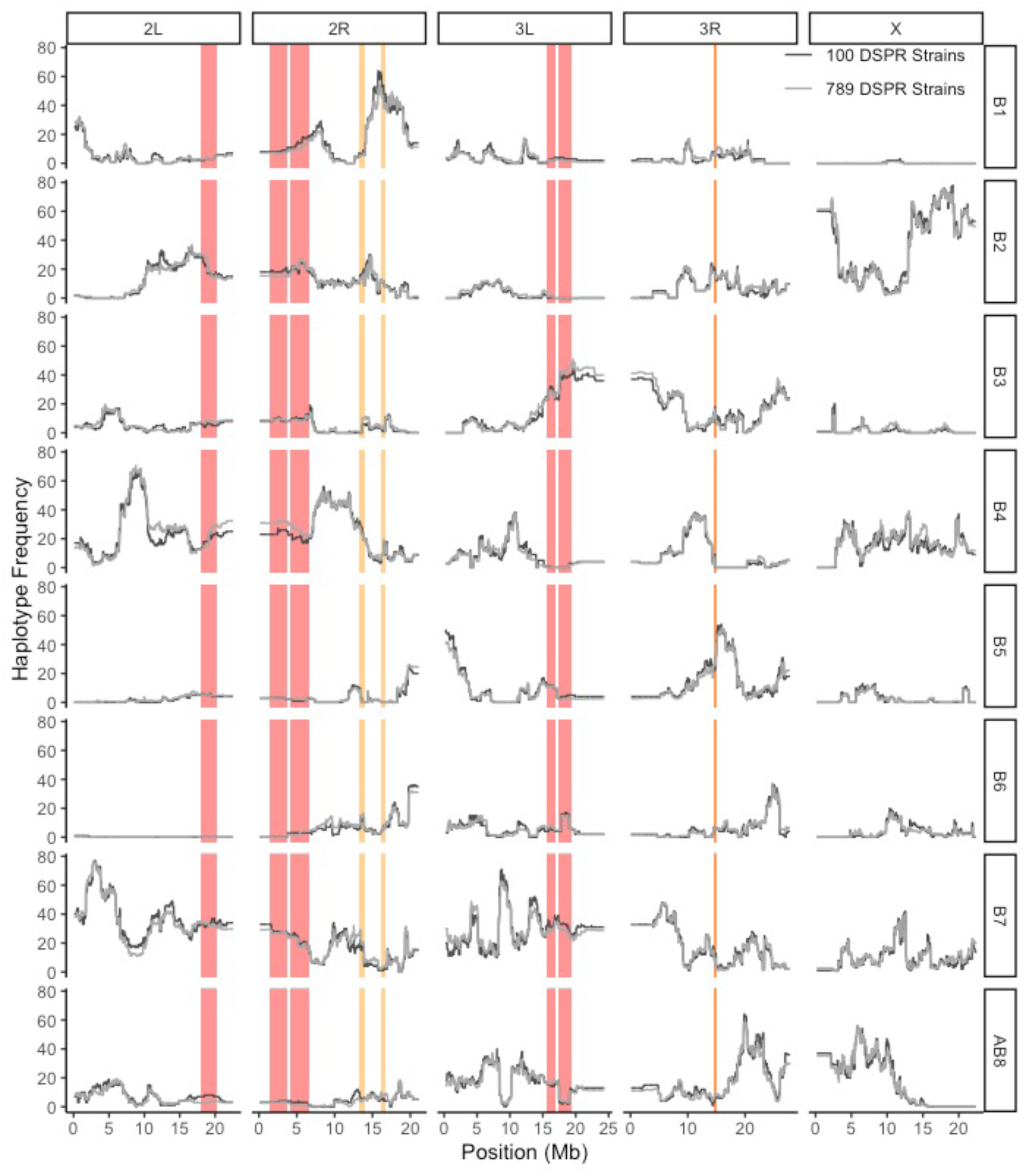
Founder haplotype frequencies are shown at each marker position (every 10,000 bp) through the genome for the 789 DSPR strains sampled for the adult copper resistance phenotype (light grey) and the 100 DSPR strains sampled for the copper-specific developmental viability phenotype (dark grey). Representation of founder haplotypes in the DSPR strains sampled for the developmental phenotype is similar to founder haplotype representation in the 789 strains sampled for the adult phenotype. In each panel, QTL intervals for adult copper resistance and copper-specific developmental viability are shown as red and yellow bars, respectively

## Supplemental Tables

**Table S1.** *Correlations between standards across each of plate and block were high and consistent*.

|  | block1_P1 | block1_P2 | block1_P3 | block2_P1 | block2_P2 | block2_P3 | block3_P1 | block3_P2 | block3_P3 |
| --- | --- | --- | --- | --- | --- | --- | --- | --- | --- |
| block1_P1 | -- | 99.99 | 99.99 | 99.97 | 99.97 | 99.91 | 99.96 | 99.94 | 99.94 |
| block1_P2 |  | -- | 99.99 | 99.98 | 99.98 | 99.90 | 99.97 | 99.96 | 99.96 |
| block1_P3 |  |  | -- | 99.98 | 99.98 | 99.92 | 99.97 | 99.95 | 99.96 |
| block2_P1 |  |  |  | -- | 99.99 | 99.86 | 99.99 | 99.98 | 99.97 |
| block2_P2 |  |  |  |  | -- | 99.87 | 99.98 | 99.97 | 99.97 |
| block2_P3 |  |  |  |  |  | -- | 99.84 | 99.86 | 99.88 |
| block3_P1 |  |  |  |  |  |  | -- | 99.98 | 99.97 |
| block3_P2 |  |  |  |  |  |  |  | -- | 99.99 |
| block3_P3 |  |  |  |  |  |  |  |  | -- |

**Table S2.**
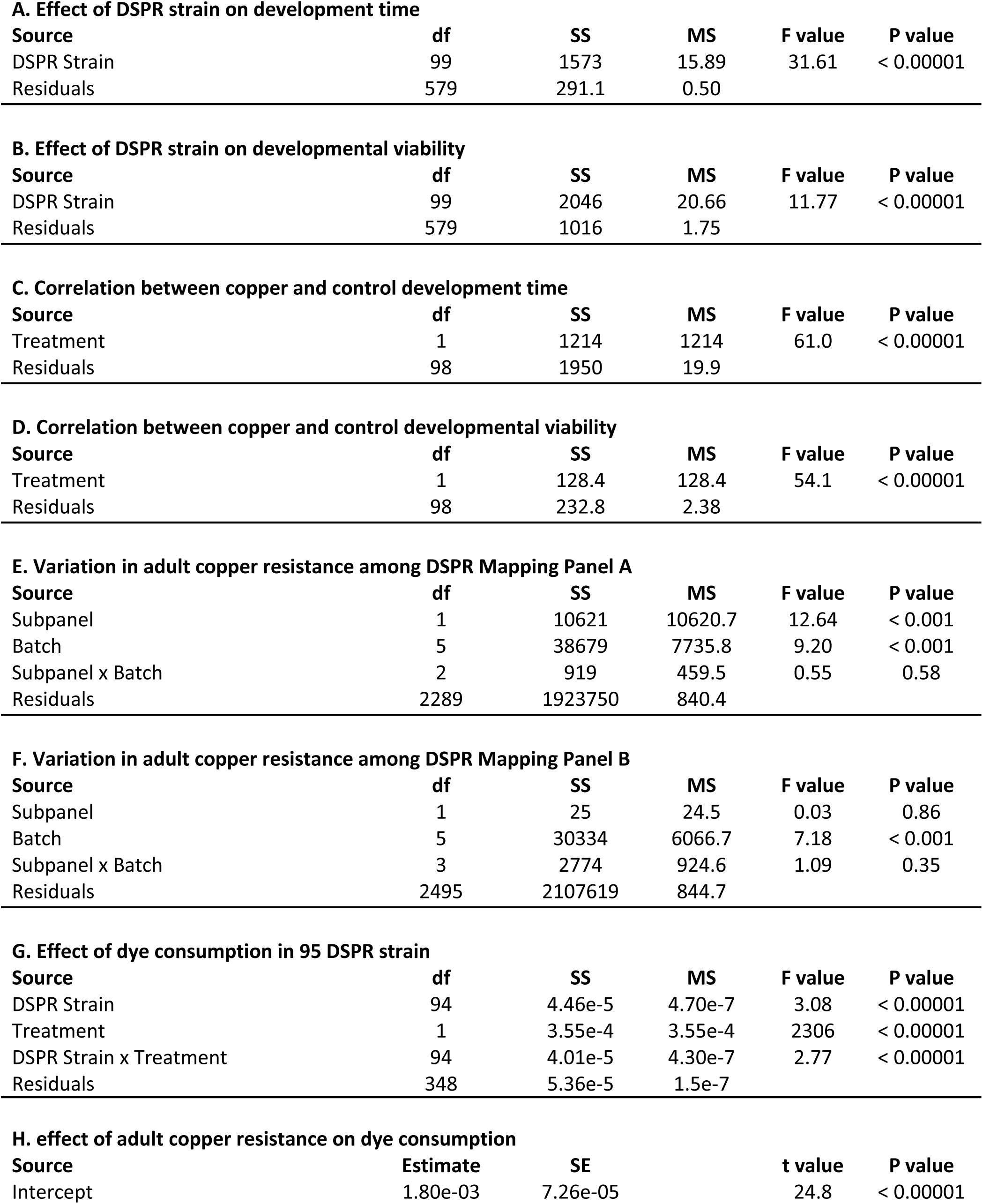
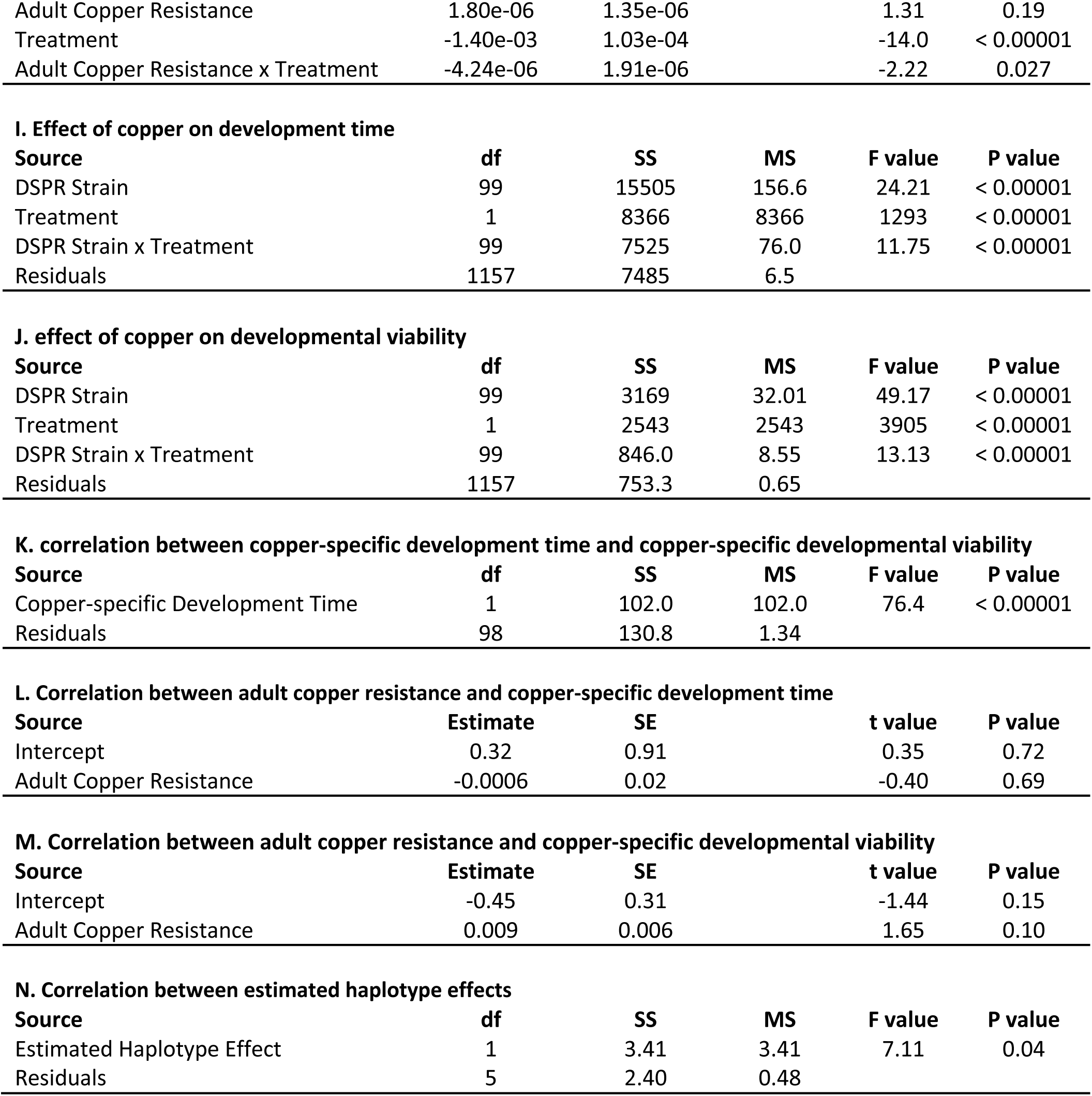
*Summary of analysis of variance and regressions*.

| **A. Effect of DSPR strain on development time** |  |  |  |  |  |
| --- | --- | --- | --- | --- | --- |
| Source | df | SS | MS | F value | P value |
| DSPR Strain | 99 | 1573 | 15.89 | 31.61 | < 0.00001 |
| Residuals | 579 | 291.1 | 0.50 |  |  |
| **B. Effect of DSPR strain on developmental viability** |  |  |  |  |  |
| Source | df | SS | MS | F value | P value |
| DSPR Strain | 99 | 2046 | 20.66 | 11.77 | < 0.00001 |
| Residuals | 579 | 1016 | 1.75 |  |  |
| **C. Correlation between copper and control development time** |  |  |  |  |  |
| Source | df | SS | MS | F value | P value |
| Treatment | 1 | 1214 | 1214 | 61.0 | < 0.00001 |
| Residuals | 98 | 1950 | 19.9 |  |  |
| **D. Correlation between copper and control developmental viability** |  |  |  |  |  |
| Source | df | SS | MS | F value | P value |
| Treatment | 1 | 128.4 | 128.4 | 54.1 | < 0.00001 |
| Residuals | 98 | 232.8 | 2.38 |  |  |
| **E. Variation in adult copper resistance among DSPR Mapping Panel A** |  |  |  |  |  |
| Source | df | SS | MS | F value | P value |
| Subpanel | 1 | 10621 | 10620.7 | 12.64 | < 0.001 |
| Batch | 5 | 38679 | 7735.8 | 9.20 | < 0.001 |
| Subpanel x Batch | 2 | 919 | 459.5 | 0.55 | 0.58 |
| Residuals | 2289 | 1923750 | 840.4 |  |  |
| **F. Variation in adult copper resistance among DSPR Mapping Panel B** |  |  |  |  |  |
| Source | df | SS | MS | F value | P value |
| Subpanel | 1 | 25 | 24.5 | 0.03 | 0.86 |
| Batch | 5 | 30334 | 6066.7 | 7.18 | < 0.001 |
| Subpanel x Batch | 3 | 2774 | 924.6 | 1.09 | 0.35 |
| Residuals | 2495 | 2107619 | 844.7 |  |  |
| **G. Effect of dye consumption in 95 DSPR strain** |  |  |  |  |  |
| Source | df | SS | MS | F value | P value |
| DSPR Strain | 94 | 4.46e-5 | 4.70e-7 | 3.08 | < 0.00001 |
| Treatment | 1 | 3.55e-4 | 3.55e-4 | 2306 | < 0.00001 |
| DSPR Strain x Treatment | 94 | 4.01e-5 | 4.30e-7 | 2.77 | < 0.00001 |
| Residuals | 348 | 5.36e-5 | 1.5e-7 |  |  |
| **H. effect of adult copper resistance on dye consumption** |  |  |  |  |  |
| Source | Estimate | SE | t value | P value |  |
| Intercept | 1.80e-03 | 7.26e-05 | 24.8 | < 0.00001 |  |

|  |  |  |  |  |
| --- | --- | --- | --- | --- |
| Adult Copper Resistance | 1.80e-06 | 1.35e-06 | 1.31 | 0.19 |
| Treatment | -1.40e-03 | 1.03e-04 | -14.0 | < 0.00001 |
| Adult Copper Resistance x Treatment | -4.24e-06 | 1.91e-06 | -2.22 | 0.027 |

I. Effect of copper on development time
| Source | df | SS | MS | F value | P value |
| --- | --- | --- | --- | --- | --- |
| DSPR Strain | 99 | 15505 | 156.6 | 24.21 | < 0.00001 |
| Treatment | 1 | 8366 | 8366 | 1293 | < 0.00001 |
| DSPR Strain x Treatment | 99 | 7525 | 76.0 | 11.75 | < 0.00001 |
| Residuals | 1157 | 7485 | 6.5 |  |  |

J. effect of copper on developmental viability
| Source | df | SS | MS | F value | P value |
| --- | --- | --- | --- | --- | --- |
| DSPR Strain | 99 | 3169 | 32.01 | 49.17 | < 0.00001 |
| Treatment | 1 | 2543 | 2543 | 3905 | < 0.00001 |
| DSPR Strain x Treatment | 99 | 846.0 | 8.55 | 13.13 | < 0.00001 |
| Residuals | 1157 | 753.3 | 0.65 |  |  |

K. correlation between copper-specific development time and copper-specific developmental viability
| Source | df | SS | MS | F value | P value |
| --- | --- | --- | --- | --- | --- |
| Copper-specific Development Time | 1 | 102.0 | 102.0 | 76.4 | < 0.00001 |
| Residuals | 98 | 130.8 | 1.34 |  |  |

L. Correlation between adult copper resistance and copper-specific development time
| Source | Estimate | SE | t value | P value |
| --- | --- | --- | --- | --- |
| Intercept | 0.32 | 0.91 | 0.35 | 0.72 |
| Adult Copper Resistance | -0.0006 | 0.02 | -0.40 | 0.69 |

M. Correlation between adult copper resistance and copper-specific developmental viability
| Source | Estimate | SE | t value | P value |
| --- | --- | --- | --- | --- |
| Intercept | -0.45 | 0.31 | -1.44 | 0.15 |
| Adult Copper Resistance | 0.009 | 0.006 | 1.65 | 0.10 |

N. Correlation between estimated haplotype effects
| Source | df | SS | MS | F value | P value |
| --- | --- | --- | --- | --- | --- |
| Estimated Haplotype Effect | 1 | 3.41 | 3.41 | 7.11 | 0.04 |
| Residuals | 5 | 2.40 | 0.48 |  |  |

**Table S3.** RNAi stocks for candidate genes.

| Gene | Stock ID | Association in This Study | Proposed Metal Association | Driver | N |
| --- | --- | --- | --- | --- | --- |
| *trpl* | RRID:BDSC_26722 | Q5B, RNAseq | Mn (Gaudet et al. 2011) | Ubiquitous | 17 |
|  |  |  |  | Midgut | 18 |
| *CG5235* | RRID:BDSC_27694 | Q8B, RNAseq | Cu (Gaudet et al. 2011) | Ubiquitous | 18 |
|  |  |  |  | Midgut | 18 |
|  | RRID:BDSC_66964 |  |  | Ubiquitous | 11 |
|  |  |  |  | Midgut | 18 |
| *ZnT41F* | RRID:BDSC_65382 | Q4B | Zn (Lye et al. 2013) | Ubiquitous | 11 |
|  |  |  |  | Midgut | 16 |
|  | RRID:BDSC_28638 |  |  | Ubiquitous | 17 |
|  |  |  |  | Midgut | 18 |
| *MtnC* | RRID:BDSC_53292 | RNAseq | Cu, Zn, Cd (Egli et al. 2006a; Calap-Quintana et al. 2017) | Ubiquitous | NA |
|  |  |  |  | Midgut | 17 |
|  | RRID:BDSC_63008 |  |  | Ubiquitous | 12 |
|  |  |  |  | Midgut | 18 |
| *Catsup* | RRID:BDSC_55396 | Q3A | Zn (Lye et al. 2013) | Ubiquitous | 4 |
|  |  |  |  | Midgut | 18 |
| *CG11825* | RRID:BDSC_58199 | Q5B | Cu (Norgate et al. 2007) | Ubiquitous | 8 |
|  |  |  |  | Midgut | 17 |
| *Ccs* | RRID:BDSC_62919 | Q5B | Cu, Zn (Kirby et al. 2008; Gaudet et al. 2011) | Ubiquitous | 12 |
|  |  |  |  | Midgut | 18 |
| *Sod1* | RRID:BDSC_34616 | NA | Cu, Zn (Gaudet et al. 2011) | Ubiquitous | 17 |
|  |  |  |  | Midgut | 18 |
| *Sod2* | RRID:BDSC_32496 |  |  | Ubiquitous | 17 |
|  |  |  |  | Midgut | 18 |
| *whd* | RRID:BDSC_33635 | Q5B, RNAseq | Fe, Cd (Strub et al. 2008) | Ubiquitous | 18 |
|  |  |  |  | Midgut | 18 |
| *DCP2* | RRID:BDSC_34806 | Q8B | Mn (Thurmond *et al.* 2019) | Ubiquitous | 17 |
|  |  |  |  | Midgut | 18 |
| *CG10505* | RRID:BDSC_38317 | RNAseq | Cu, Zn, Cd (Yepiskoposyan *et al.* 2006; Thurmond *et al.* 2019) | Ubiquitous | 16 |
|  |  |  |  | Midgut | 18 |
| *babo* | RRID:BDSC_40866 | Q5B | (Thurmond *et al.* 2019) | Ubiquitous | 15 |
|  |  |  |  | Midgut | 18 |
| *swm* | RRID:BDSC_52935 | Q3B | (Thurmond *et al.* 2019) | Ubiquitous | NA |
|  |  |  |  | Midgut | 13 |
| *Mvl* | RRID:BDSC_55316 | RNAseq | Fe, Cu, Mn, Cd (Southon et al. 2008; Bettedi et al. 2011) | Ubiquitous | 14 |
|  |  |  |  | Midgut | 18 |
| *stl* | RRID:BDSC_57811 | Q7A, RNAseq | Zn (Ozdowski et al. 2009) | Ubiquitous | 18 |
|  |  |  |  | Midgut | 18 |

**Table S4.** *Genes mapped to regions associated with each QTL. Data from FlyBase release FB2020_01. Grey text indicates non-protein coding genes. Red text indicates genes that overlap between QTL intervals*.

**Table S5.** DE genes identified with the resistance class model (∼ TRT + RES vs. reduced model: ∼ TRT), treatment model (∼ TRT + RES vs. reduced model: ∼ RES), and the full model (∼ TRT + RES vs reduced model: ∼ 1). Gene position data is from FlyBase release FB2020_01.

**Table S6.** GO terms and associated gene IDs identified for the DE genes from the full model (∼ TRT + RES vs reduced model: ∼ 1), treatment model (∼ TRT + RES vs. reduced model: ∼ RES), resistance model (∼ TRT + RES vs. reduced model: ∼ TRT), and the clusters formed for the treatment and resistance sets of DE genes. GO analysis was performed using FlyMine.

## Supplemental Files

File S1. README file for datafiles accompanying this study.

## References

1. Abu-Jamous B., and S. Kelly, 2018 Clust: Automatic extraction of optimal co-expressed gene clusters from gene expression data. https://doi.org/10.1101/221309

2. Allen W. R., 1971 Copper tolerance in some Californian populations of the monkey flower, *Mimulus guttatus*. Proc. R. Soc. Lond. B Biol. Sci. 177: 177–196. https://doi.org/10.1098/rspb.1971.0022

3. Arnold B. J., B. Lahner, J. M. DaCosta, C. M. Weisman, J. D. Hollister, et al., 2016 Borrowed alleles and convergence in serpentine adaptation. Proc. Natl. Acad. Sci. 113: 8320–8325. https://doi.org/10.1073/pnas.1600405113

4. Arya G. H., M. M. Magwire, W. Huang, Y. L. Serrano-Negron, T. F. C. Mackay, et al., 2015 The genetic basis for variation in olfactory behavior in *Drosophila melanogaster*. Chem. Senses 40: 233–243. https://doi.org/10.1093/chemse/bjv001

5. Babin-Fenske J., and M. Anand, 2011 Patterns of insect communities along a stress gradient following decommissioning of a Cu–Ni smelter. Environ. Pollut. 159: 3036–3043. https://doi.org/10.1016/j.envpol.2011.04.011

6. Bahadorani S., and A. J. Hilliker, 2009 Biological and behavioral effects of heavy metals in *Drosophila melanogaster* adults and larvae. J. Insect Behav. 22: 399–411. https://doi.org/10.1007/s10905-009-9181-4

7. Bahadorani S., S. Mukai, D. Egli, and A. J. Hilliker, 2010 Overexpression of metal-responsive transcription factor (*MTF-1*) in *Drosophila melanogaster* ameliorates life-span reductions associated with oxidative stress and metal toxicity. Neurobiol. Aging 31: 1215–1226. https://doi.org/10.1016/j.neurobiolaging.2008.08.001

8. Balamurugan K., D. Egli, A. Selvaraj, B. Zhang, O. Georgiev, et al., 2004 Metal-responsive transcription factor (*MTF-1*) and heavy metal stress response in *Drosophila* and mammalian cells: a functional comparison. Biol. Chem. 385. https://doi.org/10.1515/BC.2004.074

9. Balamurugan K., D. Egli, H. Hua, R. Rajaram, G. Seisenbacher, et al., 2007 Copper homeostasis in *Drosophila* by complex interplay of import, storage and behavioral avoidance. EMBO J. 26: 1035–1044. https://doi.org/10.1038/sj.emboj.7601543

10. Bálint A. F., M. S. Röder, R. Hell, G. Galiba, and A. Börner, 2007 Mapping of QTLs affecting copper tolerance and the Cu, Fe, Mn and Zn contents in the shoots of wheat seedlings. Biol. Plant. 51: 129–134. https://doi.org/10.1007/s10535-007-0025-9

11. Bazzicalupo A. L., J. Ruytinx, Y.-H. Ke, L. Coninx, J. V. Colpaert, et al., 2019 Incipient local adaptation in a fungus: evolution of heavy metal tolerance through allelic and copy-number variation. bioRxiv 832089; doi: https://doi.org/10.1101/832089

12. Beavis W. D., D. Grant, and R. Fincher, 1991 Quantitative trait loci for plant height in four maize populations and their associations with qualitative genetic loci. Theor. Appl. Genet. 83: 141–145.

13. Bellion M., M. Courbot, C. Jacob, F. Guinet, D. Blaudez, et al., 2007 Metal induction of a *Paxillus involutus* metallothionein and its heterologous expression in *Hebeloma cylindrosporum*. New Phytol. 174: 151–158. https://doi.org/10.1111/j.1469-8137.2007.01973.x

14. Bettedi L., M. F. Aslam, J. Szular, K. Mandilaras, and F. Missirlis, 2011 Iron depletion in the intestines of *Malvolio* mutant flies does not occur in the absence of a multicopper oxidase. J. Exp. Biol. 214: 971–978. https://doi.org/10.1242/jeb.051664

15. Blackney M. J., R. Cox, D. Shepherd, and J. D. Parker, 2014 Cloning and expression analysis of Drosophila extracellular Cu Zn superoxide dismutase. Biosci. Rep. 34: e00164. https://doi.org/10.1042/BSR20140133

16. Boyle E. A., Y. I. Li, and J. K. Pritchard, 2017 An expanded view of complex traits: from polygenic to omnigenic. Cell 169: 1177–1186. https://doi.org/10.1016/j.cell.2017.05.038

17. Bray N. L., H. Pimentel, P. Melsted, and L. Pachter, 2016 Near-optimal probabilistic RNA-seq quantification. Nat. Biotechnol. 34: 525–527. https://doi.org/10.1038/nbt.3519

18. Calap-Quintana P., J. González-Fernández, N. Sebastiá-Ortega, J. Llorens, and M. Moltó, 2017 *Drosophila melanogaster* models of metal-related human diseases and metal toxicity. Int. J. Mol. Sci. 18: 1456. https://doi.org/10.3390/ijms18071456

19. Chen S., Y. Zhou, Y. Chen, and J. Gu, 2018 fastp: an ultra-fast all-in-one FASTQ preprocessor. bioRxiv. https://doi.org/10.1101/274100

20. Cohen J., 1988 Statistical power for the behavioral sciences. Lawrence Erlbaum, Hillsdale, NJ. Collet J., and S. Fellous, 2019 Do traits separated by metamorphosis evolve independently? Concepts and methods. Proc. R. Soc. B Biol. Sci. 286: 20190445. https://doi.org/10.1098/rspb.2019.0445

21. Courbot M., G. Willems, P. Motte, S. Arvidsson, N. Roosens, et al., 2007 A major quantitative trait locus for cadmium tolerance in *Arabidopsis halleri* colocalizes with *HMA4*, a gene encoding a heavy metal ATPase. Plant Physiol. 144: 1052–1065. https://doi.org/10.1104/pp.106.095133

22. Culotta V. C., L. W. J. Klomp, J. Strain, R. L. B. Casareno, B. Krems, et al., 1997 The copper chaperone for superoxide dismutase. J. Biol. Chem. 272: 23469–23472. https://doi.org/10.1074/jbc.272.38.23469

23. Danks D. M., 1988 Copper deficiency in humans. Annu. Rev. Nutr. 8: 235–257. https://doi.org/10.1146/annurev.nu.08.070188.001315

24. Domingo A., J. González-Jurado, M. Maroto, C. Díaz, J. Vínos, et al., 1998 Troponin-T is a calcium-binding protein in insect muscle: in vivo phosphorylation, muscle-specific isoforms and developmental profile in *Drosophila melanogaster*. J. Muscle Res. Cell Motil. 19: 393–403.

25. Ecke F., N. J. Singh, J. M. Arnemo, A. Bignert, B. Helander, et al., 2017 Sublethal lead exposure alters movement behavior in free-ranging golden eagles. Environ. Sci. Technol. 51: 5729–5736. https://doi.org/10.1021/acs.est.6b06024

26. Egli D., A. Selvaraj, H. Yepiskoposyan, B. Zhang, E. Hafen, et al., 2003 Knockout of ‘metal-responsive transcription factor’ *MTF-1* in *Drosophila* by homologous recombination reveals its central role in heavy metal homeostasis. EMBO J. 22: 100–108. https://doi.org/10.1093/emboj/cdg012

27. Egli D., H. Yepiskoposyan, A. Selvaraj, K. Balamurugan, R. Rajaram, et al., 2006a A family knockout of all four *Drosophila* metallothioneins reveals a central role in copper homeostasis and detoxification. Mol. Cell. Biol. 26: 2286–2296. https://doi.org/10.1128/MCB.26.6.2286-2296.2006

28. Egli D., J. Domènech, A. Selvaraj, K. Balamurugan, H. Hua, et al., 2006b The four members of the *Drosophila* metallothionein family exhibit distinct yet overlapping roles in heavy metal homeostasis and detoxification. Genes Cells 11: 647–658. https://doi.org/10.1111/j.1365-2443.2006.00971.x

29. Ehrenreich I. M., N. Torabi, Y. Jia, J. Kent, S. Martis, et al., 2010 Dissection of genetically complex traits with extremely large pools of yeast segregants. Nature 464: 1039–1042. https://doi.org/10.1038/nature08923

30. Evans K. S., S. C. Brady, J. S. Bloom, R. E. Tanny, D. E. Cook, et al., 2018 Shared genomic regions underlie natural variation in diverse toxin responses. Genetics 210: 1509–1525. https://doi.org/10.1534/genetics.118.301311

31. Everman E. R., C. L. McNeil, J. L. Hackett, C. L. Bain, and S. J. Macdonald, 2019 Dissection of complex, fitness-related traits in multiple *Drosophila* mapping populations offers insight into the genetic control of stress resistance. Genetics 211: 19. https://doi.org/10.1534/genetics.119.301930

32. Filshie B. K., D. F. Poulson, and D. F. Waterhouse, 1971 Ultrastructure of the copper accumulating region of the *Drosophila* larval midgut. Tissue Cell 3: 77–102.

33. Freda P. J., J. T. Alex, T. J. Morgan, and G. J. Ragland, 2017 Genetic decoupling of thermal hardiness across metamorphosis in *Drosophila melanogaster*. Integr. Comp. Biol. 57: 999–1009. https://doi.org/10.1093/icb/icx102

34. Freda P. J., Z. M. Ali, N. Heter, G. J. Ragland, and T. J. Morgan, 2019 Stage-specific genotype-by-environment interactions for cold and heat hardiness in *Drosophila melanogaster*. Heredity 123: 479–491. https://doi.org/10.1038/s41437-019-0236-9

35. Gall J. E., R. S. Boyd, and N. Rajakaruna, 2015 Transfer of heavy metals through terrestrial food webs: a review. Environ. Monit. Assess. 187. https://doi.org/10.1007/s10661-015-4436-3

36. Garlapow M. E., W. Huang, M. T. Yarboro, K. R. Peterson, and T. F. C. Mackay, 2015 Quantitative genetics of food Intake in Drosophila melanogaster. PLoS ONE 10. https://doi.org/10.1371/journal.pone.0138129

37. Gaudet P., M. Livstone, and P. Thomas, 2011 Gene Ontology annotation inferences using phylogenetic trees. Go Ref. Genome Proj.

38. Georgieva S. S., S. P. McGrath, D. J. Hooper, and B. S. Chambers, 2002 Nematode communities under stress: the long-term effects of heavy metals in soil treated with sewage sludge. Appl. Soil Ecol. 20: 27–42. https://doi.org/10.1016/S0929-1393(02)00005-7

39. Goldsbrough P., 2000 Metal tolerance in plants: The role of phytochelatins and metallothioneins, pp. 227–239 in Phytoremediation of Contaminated Soil and Water, CRC Press LLC, Boca Raton, Florida.

40. GTEx Consortium, E. R. Gamazon, A. V. Segrè, M. van de Bunt, X. Wen, et al., 2018 Using an atlas of gene regulation across 44 human tissues to inform complex disease- and trait-associated variation. Nat. Genet. 50: 956–967. https://doi.org/10.1038/s41588-018-0154-4

41. Guo Y., K. Smith, J. Lee, D. J. Thiele, and M. J. Petris, 2004 Identification of methionine-rich clusters that regulate copper-stimulated endocytosis of the human Ctr1 copper transporter. J. Biol. Chem. 279: 17428–17433. https://doi.org/10.1074/jbc.M401493200

42. Guruharsha K. G., J.-F. Rual, B. Zhai, J. Mintseris, P. Vaidya, et al., 2011 A protein complex network of *Drosophila melanogaster*. Cell 147: 690–703. https://doi.org/10.1016/j.cell.2011.08.047

43. Hart E. B., H. Steenbock, J. Waddell, and C. A. Elvehjeim, 1928 Iron nutrition. VII copper is a supplement to iron for hemoglobin building in rat. J. Biol. Chem. 77: 797–812.

44. Harvey P. J., H. K. Handley, and M. P. Taylor, 2016 Widespread copper and lead contamination of household drinking water, New South Wales, Australia. Environ. Res. 151: 275–285. https://doi.org/10.1016/j.envres.2016.07.041

45. Hatori Y., and S. Lutsenko, 2013 An expanding range of functions for the copper chaperone/antioxidant protein Atox1. Antioxid. Redox Signal. 19: 945–957. https://doi.org/10.1089/ars.2012.5086

46. He X., S. Zhou, G. E. St. Armour, T. F. C. Mackay, and R. R. H. Anholt, 2016 Epistatic partners of neurogenic genes modulate *Drosophila* olfactory behavior: Epistatic modifiers for *Drosophila* olfaction. Genes Brain Behav. 15: 280–290. https://doi.org/10.1111/gbb.12279

47. Highfill C. A., G. A. Reeves, and S. J. Macdonald, 2016 Genetic analysis of variation in lifespan using a multiparental advanced intercross Drosophila mapping population. BMC Genet. 17. https://doi.org/10.1186/s12863-016-0419-9

48. Highfill C. A., B. M. Baker, S. D. Stevens, R. R. H. Anholt, and T. F. C. Mackay, 2019 Genetics of cocaine and methamphetamine consumption and preference in Drosophila melanogaster, (D. G. Heckel, Ed.). PLOS Genet. 15: e1007834. https://doi.org/10.1371/journal.pgen.1007834

49. Huard D. J. E., K. M. Kane, and F. A. Tezcan, 2013 Re-engineering protein interfaces yields copper-inducible ferritin cage assembly. Nat. Chem. Biol. 9: 169–176. https://doi.org/10.1038/nchembio.1163

50. Ilunga Kabeya F., P. Pongrac, B. Lange, M.-P. Faucon, J. T. van Elteren, et al., 2018 Tolerance and accumulation of cobalt in three species of *Haumaniastrum* and the influence of copper. Environ. Exp. Bot. 149: 27–33. https://doi.org/10.1016/j.envexpbot.2018.01.018

51. Janssens T. K. S., D. Roelofs, and N. M. van Straalen, 2009 Molecular mechanisms of heavy metal tolerance and evolution in invertebrates. Insect Sci. 16:<otherinfo> 3</otherinfo>–18. https://doi.org/10.1111/j.1744-7917.2009.00249.x

52. King E. G., S. J. Macdonald, and A. D. Long, 2012a Properties and power of the *Drosophila* Synthetic Population Resource for the routine dissection of complex traits. Genetics 191: 935–949. https://doi.org/10.1534/genetics.112.138537

53. King E. G., C. M. Merkes, C. L. McNeil, S. R. Hoofer, S. Sen, et al., 2012b Genetic dissection of a model complex trait using the *Drosophila* Synthetic Population Resource. Genome Res. 22: 1558–1566. https://doi.org/10.1101/gr.134031.111

54. King E. G., and A. D. Long, 2017 The Beavis effect in next-generation mapping panels in *Drosophila melanogaster*. G3 Genes Genomes Genet. 7: 1643–1652.

55. Kirby K., L. T. Jensen, J. Binnington, A. J. Hilliker, J. Ulloa, et al., 2008 Instability of superoxide dismutase 1 of *Drosophila* in mutants deficient for its cognate copper chaperone. J. Biol. Chem. 283: 35393–35401. https://doi.org/10.1074/jbc.M807131200

56. Lüdecke D., 2020 _sjstats: Statistical functions for regression models (Version 0.17.9). URL: https://doi.org/10.5281/zenodo.1284472

57. Lye J. C., C. D. Richards, K. Dechen, C. G. Warr, and R. Burke, 2013 In vivo zinc toxicity phenotypes provide a sensitized background that suggests zinc transport activities for most of the *Drosophila Zip* and *ZnT* genes. JBIC J. Biol. Inorg. Chem. 18: 323–332. https://doi.org/10.1007/s00775-013-0976-6

58. Lynch M., and B. Walsh, 1998 Genetic and Analysis of Quantitative Traits. Sinauer, Sunderland, MA.

59. Lyne R., R. Smith, K. Rutherford, M. Wakeling, A. Varley, et al., 2007 FlyMine: an integrated database for Drosophila and Anopheles genomics. Genome Biol. 8: R129. https://doi.org/10.1186/gb-2007-8-7-r129

60. Mackay T. F. C., S. Richards, E. A. Stone, A. Barbadilla, J. F. Ayroles, et al., 2012 The *Drosophila melanogaster* Genetic Reference Panel. Nature 482: 173–178. https://doi.org/10.1038/nature10811

61. Macnair M. R., 1983 The genetic control of copper tolerance in the yellow monkey flower, *Mimulus guttatus*. Heredity 53: 283–293.

62. Macnair M. R., 1993 The genetics of metal tolerance in vascular plants. New Phytol. 124: 541– 559. https://doi.org/10.1111/j.1469-8137.1993.tb03846.x

63. MacNair M. R., S. E. Smith, and Q. J. Cumbes, 1993 Heritability and distribution of variation in degree of copper tolerance in *Mimulus guttatus* at Copperopolis, California. Heredity 71: 445–455. https://doi.org/10.1038/hdy.1993.162

64. Maroni G., and D. Watson, 1985 Uptake and binding of cadmium, copper and zinc by *Drosophila melanogaster* larvae. Insect Biochem. 15: 55–63.

65. Marriage T. N., E. G. King, A. D. Long, and S. J. Macdonald, 2014 Fine-mapping nicotine resistance loci in *Drosophila* using a multiparent advanced generation inter-cross population. Genetics 198: 45–57. https://doi.org/10.1534/genetics.114.162107

66. Martchenko M., S. I. Candille, H. Tang, and S. N. Cohen, 2012 Human genetic variation altering anthrax toxin sensitivity. Proc. Natl. Acad. Sci. 109: 2972–2977. https://doi.org/10.1073/pnas.1121006109

67. Mercer S. W., and R. Burke, 2016 Evidence for a role for the putative *Drosophila hGRX1* orthologue in copper homeostasis. BioMetals 29: 705–713. https://doi.org/10.1007/s10534-016-9946-0

68. Mokdad R., A. Debec, and M. Wegnez, 1987 Metallothionein genes in *Drosophila melanogaster* constitute a dual system. Proc. Natl. Acad. Sci. 84: 2658–2662.

69. Montgomery S. L., D. Vorojeikina, W. Huang, T. F. C. Mackay, R. R. H. Anholt, et al., 2014 Genome-wide association analysis of tolerance to methylmercury toxicity in *Drosophila* implicates myogenic and neuromuscular developmental pathways. PLOS ONE 9: 15.

70. Moran N. A., 1994 Adaptation and constraint in the complex life cycles of animals. Annu. Rev. Ecol. Syst. 25: 573–600.

71. Najarro M. A., J. L. Hackett, and S. J. Macdonald, 2017 Loci contributing to boric acid toxicity in two reference populations of *Drosophila melanogaster*. G3 Genes Genomes Genet. 7: 1631–1641.

72. Navarro J. A., and S. Schneuwly, 2017 Copper and zinc homeostasis: lessons from Drosophila melanogaster. Front. Genet. 8. https://doi.org/10.3389/fgene.2017.00223

73. Neuberger J. S., M. Mulhall, M. C. Pomatto, J. Sheverbush, and R. S. Hassanein, 1990 Health problems in Galena, Kansas: a heavy metal mining Superfund site. Sci. Total Environ. 94: 261–272.

74. Norgate M., E. Lee, A. Southon, A. Farlow, P. Batterham, et al., 2006 Essential roles in development and pigmentation for the *Drosophila* copper transporter *DmATP7*. Mol. Biol. Cell 17: 475–484. https://doi.org/10.1091/mbc.E05-06-0492

75. Norgate M., A. Southon, S. Zou, M. Zhan, Y. Sun, et al., 2007 Copper homeostasis gene discovery in *Drosophila melanogaster*. BioMetals 20: 683–697. https://doi.org/10.1007/s10534-006-9075-2

76. Norgate M., A. Southon, M. Greenough, M. Cater, A. Farlow, et al., 2010 Syntaxin 5 is required for copper homeostasis in Drosophila and mammals, (J. M. Bridger, Ed.). PLoS ONE 5: e14303. https://doi.org/10.1371/journal.pone.0014303

77. Oliveira-Filho E. C. de, R. M. Lopes, and F. J. R. Paumgartten, 2004 Comparative study on the susceptibility of freshwater species to copper-based pesticides. Chemosphere 56: 369– 374. https://doi.org/10.1016/j.chemosphere.2004.04.026

78. Ozdowski E. F., Y. M. Mowery, and C. Cronmiller, 2009 *stall* encodes an ADAMTS metalloprotease and interacts genetically with *Delta* in *Drosophila* ovarian follicle formation. Genetics 183: 1027–1040. https://doi.org/10.1534/genetics.109.107367

79. Paradis E., J. Claude, and K. Strimmer, 2004 APE: analysis of phylogenetics and evolution in R language. Bioinformatics 20: 289–290.

80. Perkins L. A., L. Holderbaum, R. Tao, Y. Hu, R. Sopko, et al., 2015 The transgenic RNAi project at Harvard Medical School: resources and validation. Genetics 201: 843–852. https://doi.org/10.1534/genetics.115.180208

81. Petris M. J., K. Smith, J. Lee, and D. J. Thiele, 2003 Copper-stimulated endocytosis and degradation of the human copper transporter, hCtr1. J. Biol. Chem. 278: 9639–9646. https://doi.org/10.1074/jbc.M209455200

82. Pimentel H., N. L. Bray, S. Puente, P. Melsted, and L. Pachter, 2017 Differential analysis of RNA-seq incorporating quantification uncertainty. Nat. Methods 14: 687–690. https://doi.org/10.1038/nmeth.4324

83. Pinheiro J., D. Bates, S. DebRoy, D. Sarkar, and R Core Team, 2019 nlme: linear and nonlinear mixed effects models. R Package Version 31–142.

84. Plessl C., P. Jandrisits, R. Krachler, B. K. Keppler, and F. Jirsa, 2017 Heavy metals in the mallard *Anas platyrhynchos* from eastern Austria. Sci. Total Environ. 580: 670–676. https://doi.org/10.1016/j.scitotenv.2016.12.013

85. Pölkki M., 2016 The effects of copper exposure on life-history traits in insects. Ann. Univ. Turku. 328.

86. R Core Team, 2017 R: A language and environment for statistical computing. R Foundation for Statistical Computing, Vienna, Austria.

87. Ragland G. J., and J. G. Kingsolver, 2008 Evolution of thermotolerance in seasonal environments: the effects of annual temperature variation and life-history timing in *Wyeomyia smithii*. Evolution 62: 1345–1357. https://doi.org/10.1111/j.1558-5646.2008.00367.x

88. Ramirez M., S. Massolo, R. Frache, and J. A. Correa, 2005 Metal speciation and environmental impact on sandy beaches due to El Salvador copper mine, Chile. Mar. Pollut. Bull. 50: 62–72. https://doi.org/10.1016/j.marpolbul.2004.08.010

89. Roelofs D., L. Overhein, M. E. de Boer, T. K. S. Janssens, and N. M. van Straalen, 2006 Additive genetic variation of transcriptional regulation: metallothionein expression in the soil insect *Orchesella cincta*. Heredity 96: 85–92. https://doi.org/10.1038/sj.hdy.6800756

90. Ruden D. M., L. Chen, D. Possidente, B. Possidente, P. Rasouli, et al., 2009 Genetical toxicogenomics in *Drosophila* identifies master-modulatory loci that are regulated by developmental exposure to lead. NeuroToxicology 30: 898–914. https://doi.org/10.1016/j.neuro.2009.08.011

91. Ryabinina O., E. Subbian, and M. Iordanov, 2006 D-MEKKI, the *Drosophila* orthologue of mammalian *MEKK4*/*MTKI*, and *Hemipterous*/D-*MKK7* mediate the activation of D-JNK by cadmurm and arsenite in Schneider cells. BMC Cell Biol. 7: 7. https://doi.org/10.1186/1471-2121-7-7

92. Sánchez-Chardi A., and J. Nadal, 2007 Bioaccumulation of metals and effects of landfill pollution in small mammals. Part I. The greater white-toothed shrew, *Crocidura russula*. Chemosphere 68: 703–711. https://doi.org/10.1016/j.chemosphere.2007.01.042

93. Schmidt P. J., C. Kunst, and V. C. Culotta, 2000 Copper activation of superoxide dismutase 1 (SOD1) in Vivo: Role for protein-protein interactions with the copper chaperone for SOD1. J. Biol. Chem. 275: 33771–33776. https://doi.org/10.1074/jbc.M006254200

94. Selby J. P., and J. H. Willis, 2018 Major QTL controls adaptation to serpentine soils in *Mimulus guttatus*. Mol. Ecol. 27: 5073–5087. https://doi.org/10.1111/mec.14922

95. Shell B. C., R. E. Schmitt, K. M. Lee, J. C. Johnson, B. Y. Chung, et al., 2018 Measurement of solid food intake in *Drosophila* via consumption-excretion of a dye tracer. Sci. Rep. 8: 11536. https://doi.org/10.1038/s41598-018-29813-9

96. Southon A., R. Burke, M. Norgate, P. Batterham, and J. Camakaris, 2004 Copper homoeostasis in *Drosophila melanogaster* S2 cells. Biochem. J. 383: 303–309. https://doi.org/10.1042/BJ20040745

97. Southon A., A. Farlow, M. Norgate, R. Burke, and J. Camakaris, 2008 Malvolio is a copper transporter in *Drosophila melanogaster*. J. Exp. Biol. 211: 709–716. https://doi.org/10.1242/jeb.014159

98. Strub B. R., T. L. Parkes, S. T. Mukai, S. Bahadorani, A. B. Coulthard, et al., 2008 Mutations of the *withered* (*whd*) gene in *Drosophila melanogaster* confer hypersensitivity to oxidative stress and are lesions of the car*nitine palmitoyltransferase I* (*CPT I*) gene, (J. Bell, Ed.). Genome 51: 409–420. https://doi.org/10.1139/G08-023

99. Stuart G. W., P. F. Searle, and R. D. Palmiter, 1985 Identification of multiple metal regulatory elements in mouse metallothionein-I promoter by assaying synthetic sequences. Nature 317: 828–831. https://doi.org/10.1038/317828a0

100. Tchounwou P. B., C. Newsome, J. Williams, and K. Glass, 2008 Copper-induced cytotoxicity and transcriptional activation of stress genes in human liver carcinoma (HepG2) cells. Met. Ions Biol. Med. 10: 285–290.

101. Terhzaz S., P. Cabrero, V. R. Chintapalli, S.-A. Davies, and J. A. T. Dow, 2010 Mislocalization of mitochondria and compromised renal function and oxidative stress resistance in *Drosophila SesB* mutants. Physiol. Genomics 41: 33–41. https://doi.org/10.1152/physiolgenomics.00147.2009

102. Thurmond J., J. L. Goodman, V. B. Strelets, H. Attrill, L. S. Gramates, et al., 2019 FlyBase 2.0: the next generation. Nucleic Acids Res. 47: D759–D765. https://doi.org/10.1093/nar/gky1003

103. Turner T. L., E. C. Bourne, E. J. Von Wettberg, T. T. Hu, and S. V. Nuzhdin, 2010 Population resequencing reveals local adaptation of *Arabidopsis lyrata* to serpentine soils. Nat. Genet. 42: 260–263. https://doi.org/10.1038/ng.515

104. Turski M. L., and D. J. Thiele, 2007 *Drosophila Ctr1A* functions as a copper transporter essential for development. J. Biol. Chem. 282: 24017–24026. https://doi.org/10.1074/jbc.M703792200

105. Uriu-Adams J. Y., and C. L. Keen, 2005 Copper, oxidative stress, and human health. Mol. Aspects Med. 26: 268–298. https://doi.org/10.1016/j.mam.2005.07.015

106. Usmani J. M., 2011 Accumulation of heavy metals in fishes: a human health concern. Int. J. Environ. Sci. 2: 671–682.

107. Wang X., and L. Kruglyak, 2014 Genetic basis of haloperidol resistance in Saccharomyces cerevisiae Is complex and dose dependent, (G. P. Copenhaver, Ed.). PLoS Genet. 10: e1004894. https://doi.org/10.1371/journal.pgen.1004894

108. Willems G., D. B. Dräger, M. Courbot, C. Godé, N. Verbruggen, et al., 2007 The genetic basis of zinc tolerance in the metallophyte *Arabidopsis halleri* ssp. *halleri* (Brassicaceae): An analysis of quantitative trait loci. Genetics 176: 659–674. https://doi.org/10.1534/genetics.106.064485

109. World Health Organization, Food and Agriculture Organization of the United Nations, and International Atomic Energy Agency (Eds.), 1996 Trace elements in human nutrition and health. World Health Organization, Geneva.

110. Wright K. M., U. Hellsten, C. Xu, A. L. Jeong, A. Sreedasyam, et al., 2015 Adaptation to heavy-metal contaminated environments proceeds via selection on pre-existing genetic variation. http://biorxiv.org/lookup/doi/10.1101/029900

111. Wu L., A. D. Bradshaw, and D. A. Thurman, 1975 The potential for evolution of heavy metal tolerance in plants. III. The rapid evolution of copper tolerance in Agrostis stolonifera. Heredity 34: 165–187.

112. Wuana R. A., and F. E. Okieimen, 2011 Heavy metals in contaminated soils: a review of sources, chemistry, risks and best available strategies for remediation. ISRN Ecol. 2011: 1–20. https://doi.org/10.5402/2011/402647

113. Yepiskoposyan H., D. Egli, T. Fergestad, A. Selvaraj, C. Treiber, et al., 2006 Transcriptome response to heavy metal stress in *Drosophila* reveals a new zinc transporter that confers resistance to zinc. Nucleic Acids Res. 34: 4866–4877. https://doi.org/10.1093/nar/gkl606

114. Yin S., Q. Qin, and B. Zhou, 2017 Functional studies of *Drosophila* zinc transporters reveal the mechanism for zinc excretion in Malpighian tubules. BMC Biol. 15: 12. https://doi.org/10.1186/s12915-017-0355-9

115. Zhou H., K. M. Cadigan, and D. J. Thiele, 2003 A Copper-regulated transporter required for copper acquisition, pigmentation, and specific stages of development in *Drosophila melanogaster*. J. Biol. Chem. 278: 48210–48218. https://doi.org/10.1074/jbc.M309820200

116. Zhou S., T. V. Morozova, Y. N. Hussain, S. E. Luoma, L. McCoy, et al., 2016 The genetic basis for variation in sensitivity to lead toxicity in Drosophila melanogaster. Environ. Health Perspect. 124. https://doi.org/10.1289/ehp.1510513

117. Zhou S., S. E. Luoma, G. E. St. Armour, E. Thakkar, T. F. C. Mackay, et al., 2017 A Drosophila model for toxicogenomics: Genetic variation in susceptibility to heavy metal exposure, (G. P. Copenhaver, Ed.). PLOS Genet. 13: e1006907. https://doi.org/10.1371/journal.pgen.1006907

